# Na^+^/H^+^ antiport across the thylakoid membrane is functionally conserved in the Chloroplastida *Arabidopsis*

**DOI:** 10.64898/2026.11.22.768382

**Authors:** Eduard Windenbach, Pedro Barreto, Josha Ebeling, Quang Huy Nguyen, Geoffry A. Davis, Deserah D. Strand, Charlotte Seydel, Benjamin Brandt, Bettina Bölter, Susanne Mühlbauer, Lorenz Holzner, Beata Szulc, Andreas Klingl, Serena Schwenkert, Heinrich Jung, Ute Krämer, Markus Schwarzländer, Anna Matuszynska, Ute Armbruster, Hans-Henning Kunz

**Author notes:** Corresponding author: Hans-Henning Kunz.

## Abstract

In nature, plants experience frequently changing light conditions. To balance photoprotection and photosynthetic efficiency, chloroplasts dynamically regulate proton-motive force (PMF) partitioning through thylakoid ion transport proteins. While the roles of K^+^ and Cl^-^ transport in PMF regulation are established in *Arabidopsis thaliana*, the contribution of Na^+^ flux remains unexplored. Here, we identify NHD1 as a thylakoid-localized transporter that mediates pH-dependent Na^+^/H^+^ antiport in a heterologous system with a K_m_ of approximately 2 mM. Unlike other characterized thylakoid ion transport proteins, NHD1 loss does not alter photosynthesis when plants are grown under control light. However, prolonged fluctuating-light exposure uncovered pronounced physiological consequences of NHD1 deficiency: null mutants exhibit delayed non-photochemical quenching (NPQ) relaxation, reduced photosystem II quantum yield, altered chloroplast ultrastructure, and significantly impaired growth. Also, higher-order mutants lacking NHD1, K^+^/H^+^ exchanger KEA3, and voltage-dependent chloride channel VCCN1, exhibit stronger photosynthetic responses than the respective single mutants, suggesting partial functional redundancy among these thylakoid ion transporters. Together, our findings establish Na^+^/H^+^ exchange via NHD1 as a previously unrecognized, conserved component of thylakoid ion homeostasis, highlighting its importance for long-term photosynthetic acclimation to dynamic light environments, and expanding the repertoire of PMF-regulating transporters.

**One-sentence summary:** NHD1-mediated Na^+^/H^+^ exchange across the thylakoid membrane is conserved in Chloroplastida and required for long-term acclimation to dynamic light environments.

## Introduction

Oxygenic photosynthesis is the primary basis of most aerobic multicellular life on Earth (Schirrmeister et al., 2013). In phototrophic eukaryotes, photosynthesis takes place in the chloroplast. It can be subdivided into light-dependent reactions at the thylakoid membrane and light-independent reactions, i.e., CO_2_ fixation in the Calvin-Benson-Bassham cycle (CBBC) located in the stroma (Rochaix, 2011). Light energy drives charge separation in the photosystems, resulting in water splitting at photosystem II (PSII) and concomitant e^-^ transport from PSII via the cytochrome *b*_6_*f* complex (*b*_6_*f*) towards photosystem I (PSI) (McEvoy and Brudvig, 2006). At the stromal side of PSI, e^-^ are transferred to ferredoxin (Fd) and are subsequently used by the Fd-NADP^+^ oxidoreductase (FNR) to reduce NADP^+^ (Mulo, 2011), finalizing the so-called linear electron flow (LEF) process. During LEF, H^+^ are transferred into the thylakoid lumen, lowering its pH and setting up a proton motive force (PMF).

The PMF drives stromal ATP production by the ATP synthase (Mitchell, 1966). Over the last decade, our understanding of the multiple ways in which the PMF contributes to plant productivity has increased significantly. For a long time, ΔpH was considered the predominant component of the plastid PMF, while the contribution of the trans-thylakoid membrane potential (Δψ) was strongly underappreciated (Bulychev and Vredenberg, 1999). However, depending on the prevailing light conditions, the PMF can change in magnitude and partition into ΔpH and Δψ (Witt, 1979; Cruz et al., 2001). A high H^+^ concentration in the lumen decreases e^-^ flow through the cytochrome *b*_6_*f* complex (photosynthetic control) (Degen and Johnson, 2024) and activates energy dependent quenching (qE), i.e., regulated energy dissipation as heat, the main component of non-photochemical quenching in plants (NPQ) (Hope et al., 1994; Demmig-Adams et al., 1996). Photosynthetic control and NPQ are critical components in protecting the photosynthetic machinery from damage during light stress. However, when the light conditions change, e.g., after a sudden drop in intensity, photoprotection becomes a limiting factor for efficient photochemistry by dissipating light energy as heat even when photochemical capacity could be safely restored (Zhu et al., 2004). To fine-tune the balance between photoprotection and photosynthetic efficiency in response to changing light conditions, chloroplasts dynamically regulate PMF partitioning between ΔpH and Δψ, allowing rapid adjustment of luminal acidification (Armbruster et al., 2017). Thus, identifying the molecular components that regulate PMF *in vivo* could enable targeted strategies to accelerate NPQ relaxation under fluctuating light – a transition currently associated with a significant but avoidable loss of photochemical efficiency and, consequently, plant yield (Zhu et al., 2004; Kromdijk et al., 2016; Lehretz et al., 2022).

Thylakoid membrane ion transport proteins have attracted considerable research attention in recent years. The activity of ion channels and H^+^-dependent carriers enables a plant to rapidly switch the relative contribution of Δψ and ΔpH to PMF. Several candidates have been suggested in *Arabidopsis*, but the thylakoid ion transportome is still incomplete (Kunz et al., 2024). One of the known transporters is the K^+^/H^+^ exchanger KEA3, operating as an electroneutral antiporter that exports protons from the thylakoid lumen into the stroma in exchange for K^+^ influx. By doing so, it directly decreases the ΔpH component while leaving total PMF unaffected – effectively repartitioning PMF toward Δψ (Armbruster et al., 2014; Kunz et al., 2014). When *kea3* loss-of-function mutant plants are shifted into fluctuating light, the missing K^+^/H^+^ exchange results in delayed NPQ inactivation (Armbruster et al., 2016; Wang et al., 2017). A second protein of particular relevance to PMF regulation is the thylakoid voltage-dependent Cl^-^ channel VCCN1. By importing negatively charged Cl^-^ into the lumen, VCCN1 effectively compensates for the positive charge created by proton accumulation, thereby promoting the conversion of Δψ into ΔpH. Hence, its loss prevents rapid NPQ activation in response to dynamic light (Duan et al., 2016; Herdean et al., 2016; Li et al., 2021). Lastly, CLCe also resides in the thylakoid membrane and has been proposed to mediate Cl^-^/H^+^ exchange (Marmagne et al., 2007; Herdean et al., 2016). However, its overall impact on photosynthesis is comparatively mild relative to KEA3 and VCCN1 (Li et al., 2021). Even though the aforementioned transport proteins have been thoroughly characterized, export pathways for K^+^ and Cl^-^ from the thylakoid lumen remain unknown (Höhner et al., 2019). Furthermore, other ions with potential effects on PMF regulation have not yet been considered.

Cyanobacteria, the evolutionary predecessors of chloroplasts, exchange Na^+^ ions across their thylakoids. In the model *Synechocystis* sp. PCC 6803, the Na^+^/H^+^ antiporter NhaS3 resides in the thylakoid membrane (Tsunekawa et al., 2009; Tsujii et al., 2025), where its transport seems critical for cell viability, as chromosomal segregation of the *nhaS3* knockout could not be achieved (Inaba et al., 2001; Wang et al., 2002). The precise physiological basis for the essentiality of NhaS3 remains to be elucidated. Among all Na^+^/H^+^ antiporters encoded in the *Synechocystis* sp. PCC 6803 genome, NhaS3 is the closest homolog of the *Arabidopsis* K^+^/H^+^ antiporter KEA3 (Tsujii et al., 2019). Both proteins localize to the thylakoid membrane, yet their ion selectivity differs: NhaS3 functions as a Na^+^/H^+^ antiporter, whereas KEA3 mediates K^+^/H^+^ exchange (Tsujii et al., 2019). Whether the two transporters fulfill analogous physiological roles remains an open question. To date, no Na^+^/H^+^ transporter has been described in the thylakoid membranes of any Chloroplastida, raising the following question: Have plants lost trans-thylakoid Na^+^/H^+^ exchange, perhaps reflecting lower physiological relevance and abundance of Na^+^ relative to K^+^ or do other, yet uncharacterized, carriers mediate Na^+^/H^+^ exchange?

Interestingly, Chloroplastida genomes also encode alternative cation/H^+^ antiporters, such as D-type Na^+^/H^+^ antiporters (NhaD). NhaD proteins belong to the ion transporter (IT) superfamily (Patiño-Ruiz et al., 2022). They are primarily found in alkaliphile and halophile bacteria, mediating Na^+^(Li^+^)/H^+^ exchange (Lv et al., 2022). In *Arabidopsis*, one NhaD protein designated NHD1 (At3g19490) is present in chloroplasts. However, its exact sub-organellar localization is unclear. Initial microscopy in plants and moss suggested an association with the chloroplast envelope (Barrero-Gil et al., 2007; Cosentino et al., 2010; Müller et al., 2014), while NHD1 has also been detected in enriched thylakoid fractions of *Arabidopsis thaliana* by mass spectrometry (Tomizioli et al., 2014). If NHD1 were to localize to the thylakoid membrane, it could plausibly facilitate Na^+^/H^+^ exchange and affect PMF regulation, providing a previously unknown Na^+^-dependent regulatory mechanism within the green lineage.

In this study of the model plant *Arabidopsis thaliana*, we set out to a) determine NHD1’s exact protein localization, b) verify NHD1 transport properties, c) probe whether chloroplasts accumulate Na^+^ at concentrations sufficient to support flux through NHD1, and d) test NHD1’s relevance for photosynthetic efficiency.

## Results

### NHD1 localizes to the thylakoid membrane

As described above, the sub-organellar localization of NHD1 within plastids has remained ambiguous. Therefore, NHD1-mVenus was co-expressed with established chloroplast membrane marker proteins TIC40-mCherry, an inner envelope protein, and KEA3-mCherry, a thylakoid-localized antiporter (Höhner et al., 2019). Following transient expression in tobacco leaves, the NHD1-mVenus fluorescence matched the chlorophyll autofluorescence in both co-expression experiments (Fig. 1A, B), suggesting that NHD1 is localized to the thylakoid membrane. TIC40-mCherry exhibited the characteristic ring-like fluorescence pattern of chloroplast envelope membrane proteins, while the signal distribution of the NHD1-mVenus fusion protein was clearly distinct (Fig. 1A). In contrast, the KEA3-mCherry signal overlapped with both chlorophyll autofluorescence and the NHD1-mVenus signal (Fig. 1B), indicating that both transporters reside within the thylakoid membrane.

**Fig. 1.**
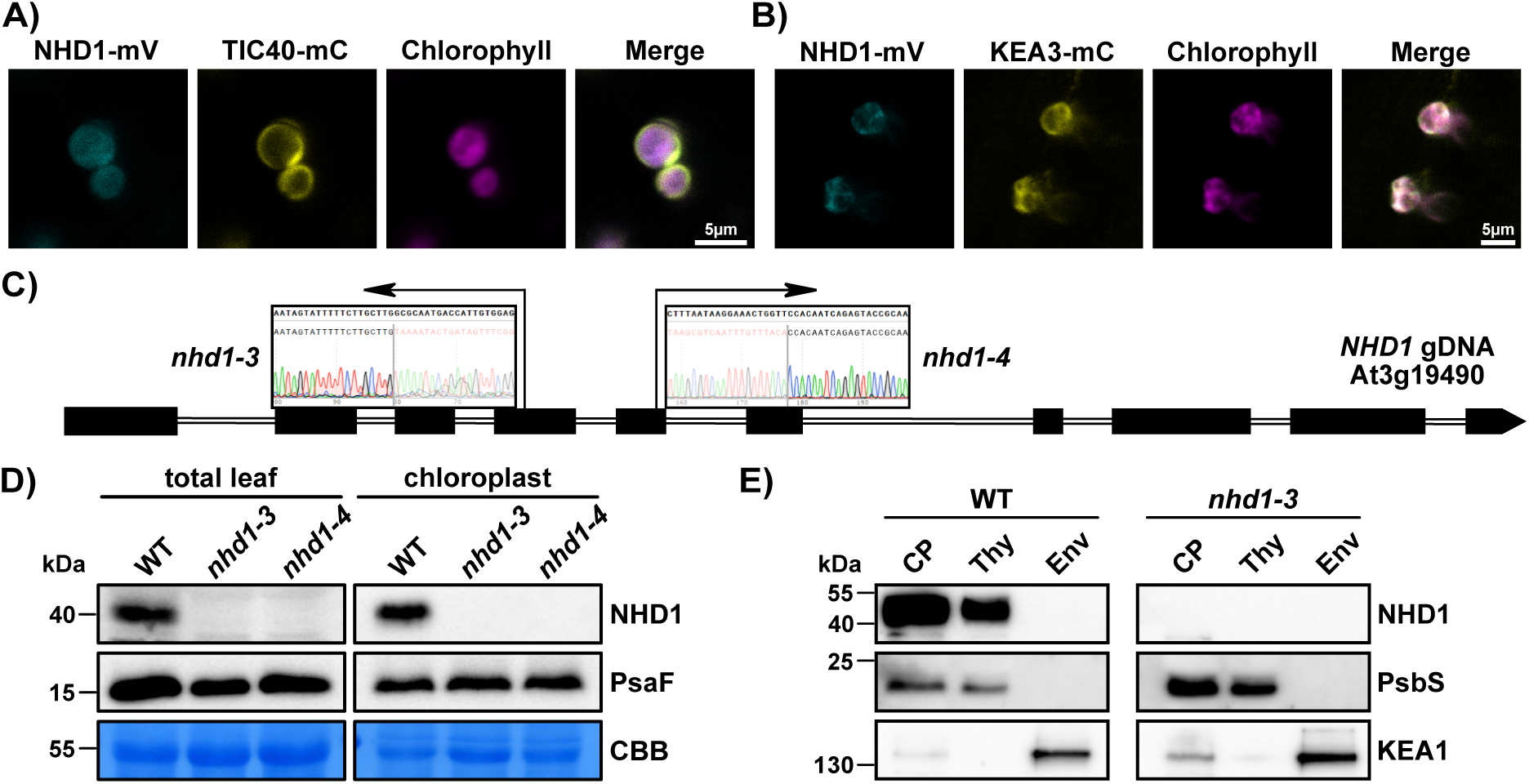
NHD1 is a thylakoid membrane protein. Confocal laser microscopy of transiently co-transformed tobacco leaf discs with A) NHD1-mVenus and plastid inner envelope marker TIC40-mCherry. NHD1-mVenus fluorescence in cyan, TIC40-mCherry fluorescence in yellow, chlorophyll autofluorescence in magenta and a merged image are depicted; Scale bar = 5 µm. B) NHD1-mVenus and thylakoid membrane marker KEA3-mCherry. NHD1-mVenus fluorescence in cyan, KEA3-mCherry fluorescence in yellow, chlorophyll autofluorescence in magenta and a merged image are depicted. Scale bar = 5 µm. C) Schematic representation of exon and intron structure of the *NHD1* locus, indicating the T-DNA insertion sites of the *nhd1-3* (exon 4) and *nhd1-4* (exon 5) alleles. D) Immunoblot analysis of NHD1 in normalized total leaf and isolated chloroplast protein extracts from WT, *nhd1-3*, and *nhd1-4*. PsaF detection and Coomassie Brilliant Blue (CBB) staining of RuBisCO LSU serve as loading control. E) Immunoblot analysis of NHD1 in isolated chloroplast and subplastidial fractions. Chloroplasts (CP), thylakoid (Thy) and envelope (Env) proteins were analyzed. Detection of PsbS (thylakoid marker) and KEA1 (envelope marker) were used as controls to probe the purity of each fraction.

To verify the sub-organellar localization, we obtained two homozygous loss-of-function alleles: *nhd1-3* (Koncz et al., 1992) and *nhd1-4* (SALK_118891) (Fig. 1C; Fig. S1A, B). The additional NHD1 mutant lines were needed since previously described *nhd1* loss-of-function mutants carry second-site insertions and large deletions (Lopez et al., 2022). We successfully confirmed the loss of NHD1 in leaf protein extracts and isolated plastids from *nhd1-3* and *nhd1-4* by immunoblot (Fig. 1D). Subsequently, we prepared envelope and thylakoid subfractions from isolated WT and *nhd1* chloroplasts. Immunoblots on the isolated fractions confirmed that NHD1 was detected in total chloroplasts and the thylakoid membrane fraction but not in the envelope membrane fraction in multiple independent experiments (Fig. 1E; Fig. S1C). The purity of membrane fractions was confirmed by the presence of K^+^/H^+^ exchanger KEA1 in the envelope and the photosystem II subunit PsbS in the thylakoid membrane, respectively (Li et al., 2000; Bölter et al., 2020).

### NHD1 shows Na^+^/H^+^ antiporter activity

Although Na^+^ movement across the thylakoid membrane was inferred early on from spectroscopic measurements of membrane potential changes (Barber, 1972), these experiments could not establish whether ion movement occurs via protein-mediated transport. The detailed kinetic characterization of a thylakoid-localized Na^+^/H^+^ antiporter would provide the first molecular basis for regulated Na^+^ transport across this membrane. Hence, we analyzed transport properties of NHD1 in a recombinant system. NHD1 was introduced into the salinity-sensitive *E. coli* strain TO114 which carries chromosomal deletions of the three major endogenous Na^+^/H^+^ antiporter-encoding genes, rendering it devoid of endogenous Na^+^/H^+^ antiport activity (Ohyama et al., 1994).

As a positive control, TO114 was separately transformed with the cyanobacterial thylakoid Na^+^/H^+^ antiporter NhaS3 from *Synechocystis*. In line with previous work (Inaba et al., 2001), NhaS3 expression rescued salt-sensitive growth on plates containing 300 mM NaCl. Similarly, NHD1 expression in TO114 complemented the salt-sensitive phenotype, indicating that the protein does exhibit Na^+^ transport activity (Fig. 2A; Fig. S2A). Given that the moss ortholog PpNHAD1 showed evidence of mediating K^+^ flux in a heterologous expression system (Barrero-Gil et al., 2007), K^+^ transport of AtNHD1 was re-examined using the K^+^ uptake-deficient *E. coli* strain LB2003 (Stumpe and Bakker, 1997). The established thylakoid K^+^/H^+^ antiporter KEA3 was used as a positive control for this experiment. As previously reported (Tsujii et al., 2019), KEA3 rescued the *E. coli* mutant, showing improved growth on synthetic medium containing only traces of K^+^. Under these conditions, however, NHD1 was unable to rescue the growth of the *E. coli* LB2003 cells to the same extent as KEA3 (Fig. 2B; Fig. S2B), suggesting that NHD1 does not mediate K^+^ transport with comparable affinity.

**Fig. 2.**
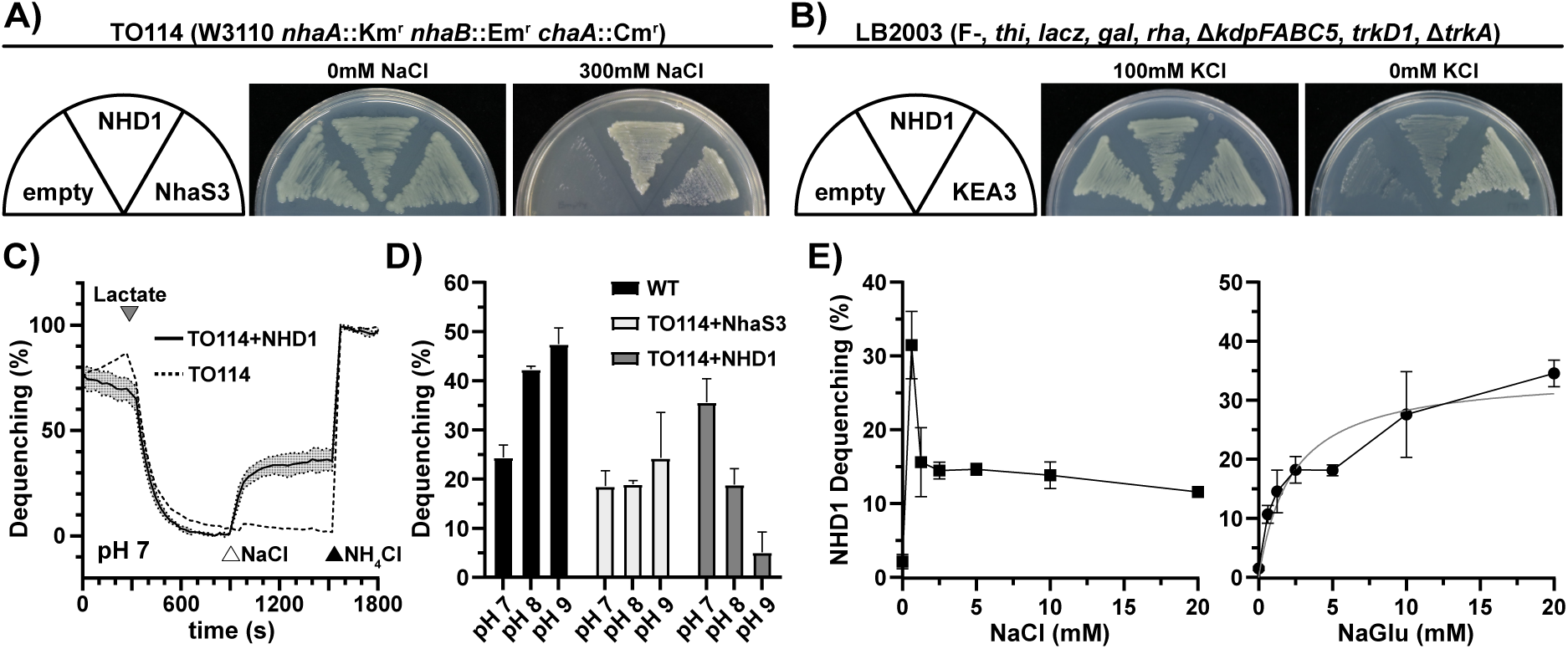
NHD1 catalyzes Na^+^/H^+^ exchange. A) Growth complementation assay of Na^+^ export-deficient *E. coli* strain (TO114) transformed with *NHD1*, *Synechocystis sp.* PCC 6803 *NhaS3* (positive control), and an empty vector (negative control). Cells were grown in the presence of 300 mM NaCl. B) Growth complementation assay of a K^+^ import deficient *E. coli* strain (LB2003) transformed with NHD1, KEA3 (positive control), and an empty vector (negative control). Cells were grown on LB0 medium and LB0 medium supplemented with 100 mM KCl. C) Na^+^/H^+^ antiporter activity of NHD1 in inverted membrane vesicles of *E. coli*. Addition of D-Lactate (grey triangle) produces a ΔpH i.e. acridine orange fluorescence is quenched. Antiporter activity is initiated by the addition of NaCl (white triangle). Lastly, ΔpH is dissipated by the addition of NH_4_Cl (black triangle). TO114 was used as a negative control (avg. ± SD, n=3). D) pH dependence of NHD1-mediated antiport in comparison to native Na^+^/H^+^ antiporters of WT and *Synechocystis sp.* PCC 6803 NhaS3 (avg. ± SD, n=3). E) Concentration dependent Na^+^/H^+^ antiport of NHD1 using NaCl (left panel) or Na-D-gluconate (NaGlu) (right panel) at pH 8.0 for NHD1 containing vesicles (avg. ± SD, n=3). The Michaelis-Menten fit is shown in gray for NaGlu induced antiport.

To corroborate these observations and characterize H^+^-dependent antiport activity in more detail, antiport assays were performed using inverted vesicles from an *E. coli* wild-type strain, as a positive control, TO114, as a negative control, and TO114 expressing either NhaS3 or NHD1. NHD1 expression was verified in the vesicle preparations by immunoblotting (Fig. S2C). Addition of lactate allowed its oxidation by membrane-bound lactate dehydrogenase and concomitant H^+^ relocation via the respiratory chain complexes into the vesicles, setting up a ΔpH (Rosen and McClees, 1974). This was monitored via fluorescence quenching of the preloaded pH dye acridine orange (Fig. 2C). Upon the addition of NaCl, vesicles prepared from wild-type cells exhibited pronounced fluorescence dequenching indicative of robust Na^+^/H^+^ antiporter activity (Fig. S2D) while those prepared from the TO114 mutant displayed only minimal dequenching, consistent with the loss of endogenous Na^+^/H^+^ antiporters (Fig. S2D). In contrast, TO114 transformed with NhaS3 showed clear Na^+^/H^+^ antiport (Fig. S2D). Na^+^/H^+^ exchange was also observed in vesicles from TO114 cells expressing NHD1 (Fig. 2C), thereby confirming its function as a Na^+^/H^+^ antiporter. Consistent with measurements of other NhaD transporters (Liu et al., 2005), NHD1 was also active with LiCl (Fig. S2E). No K^+^/H^+^ antiport activity was detected with KCl, whereas slight Rb^+^-dependent dequenching was observed under the same conditions, albeit substantially lower in magnitude than the dequenching observed with NaCl and LiCl (Fig. S2E). This discrepancy in K^+^ and Rb^+^-dependent dequenching may indicate that NHD1-mediated K^+^ transport, if present at all, is below the detection threshold of the dequenching assay.

In addition to substrate specificity, the pH dependence of transport activity was investigated. The endogenous *E. coli* antiporters displayed maximal activity under alkaline conditions (Fig. 2D), consistent with the pronounced pH dependence of the NhaA antiporter (Taglicht et al., 1991). In addition, the wild type exhibited substantial antiport activity at neutral pH (Fig. 2D), which is facilitated by NhaB, a pH-independent Na^+^/H^+^ exchanger (Pinner et al., 1994). As previously reported (Tsunekawa et al., 2009), NhaS3 maintained relatively similar activity across the entire pH range tested (Fig. 2D). In contrast to other NhaD transporters, characteristically active at alkaline pH (Lv et al., 2022), NHD1 exhibited an inverse pH dependence, with maximal dequenching observed at neutral pH and a clear activity decrease under alkaline conditions, where transport was almost completely abolished (Fig. 2D). This pronounced pH sensitivity may represent a regulatory mechanism controlling NHD1 activity *in planta*, although the molecular basis of this regulation remains to be elucidated.

To test whether the observed transport activity genuinely stems from Na^+^ and not from added Cl^-^ ions, sodium gluconate was employed. The persistent Na^+^-dependent fluorescence quenching observed for NhaS3 (Fig. S2F) and NHD1 (Fig. 2E) confirms that both carriers move Na^+^ cations and not anions across the membrane. To determine the substrate affinity of NHD1, Na^+^-dependent fluorescence dequenching was measured over a range of salt concentrations and fitted using Michaelis-Menten kinetics. For reasons that remain unclear and are beyond the scope of the present characterization, NHD1 exhibited pronounced activity at low NaCl concentrations (≤1.25 mM) (Fig. 2E, left panel). This behavior prevented a reliable fit to a Michaelis-Menten model. Therefore, kinetic analysis was performed using sodium gluconate. Here, NHD1 exhibited a K_m_ of 2.2 mM, which was comparable to that of NhaS3 (2.7 mM) (Fig. 2E, right panel; Fig. S2F) and fell within the Na^+^ affinity range reported for other members of the NhaD transporter family (Dzioba et al., 2002; Liu et al., 2005). In summary, our results identify NHD1 as a pH-regulated Na^+^/H^+^ antiporter in the thylakoid membrane. NHD1’s transport activity and substrate affinity resemble those of other characterized members of the NhaD family, while its pH dependence appears to be different.

### The functional relevance of NHD1 becomes apparent in the absence of other thylakoid transporters

To assess whether plastid Na^+^ concentrations in *Arabidopsis* are within a range sufficient for NHD1 transport activity, we determined Na^+^ levels in isolated chloroplasts (Fig. S3A). Consistent with previous reports (Müller et al., 2014), the detected amounts of approximately 5 ng Na^+^ (µg chlorophyll)^-1^ were within a range required to support flux through NHD1. In line with its localization in the thylakoid membrane, loss of NHD1 did not result in detectable alterations in chloroplast Na^+^ content (Fig. S3A). Ion transporters and channels play a crucial role in PMF dynamics and H^+^ transfer across the thylakoid membrane. Previous analyses of higher-order loss-of-function mutants demonstrated that K^+^/H^+^ antiporter KEA3 and the voltage-dependent Cl^-^ channel VCCN1 each regulate photosynthesis independently, with no evidence of functional compensation among them (Dukic et al., 2019; Li et al., 2021). However, this independence of function does not preclude the possibility that either transporter could partially compensate for the absence of an as-yet-uncharacterized transporter contributing to chloroplast proton flux. To test this, we produced higher-order loss-of-function mutants by crossing previously described thylakoid transporter/channel mutants (*kea3* and *vccn1*) with *nhd1* loss-of-function mutants (Fig. S3D-F).

Measurements with *nhd1* mutants showed only minor differences in the quantum yield of regulated non-photochemical quenching (Y(NPQ)), with a slightly elevated NPQ response in the early phase of light treatment (Fig. 3B). Changes in trans-thylakoid H^+^ and ion flux may correlate with altered PMF fractions. To investigate PMF partitioning, the electrochromic shift (ECS) of the carotenoids, a measure of the membrane potential (Δψ) across the thylakoid membrane, was probed (Witt, 1979; Cruz et al., 2001). Interestingly, while the size of both PMF components was unaltered in mutants, the ECS_inv_ kinetics, reflecting counterion movements, differed between both *nhd1* loss-of-function lines and WT controls (Fig. S3B, C).

**Fig. 3.**
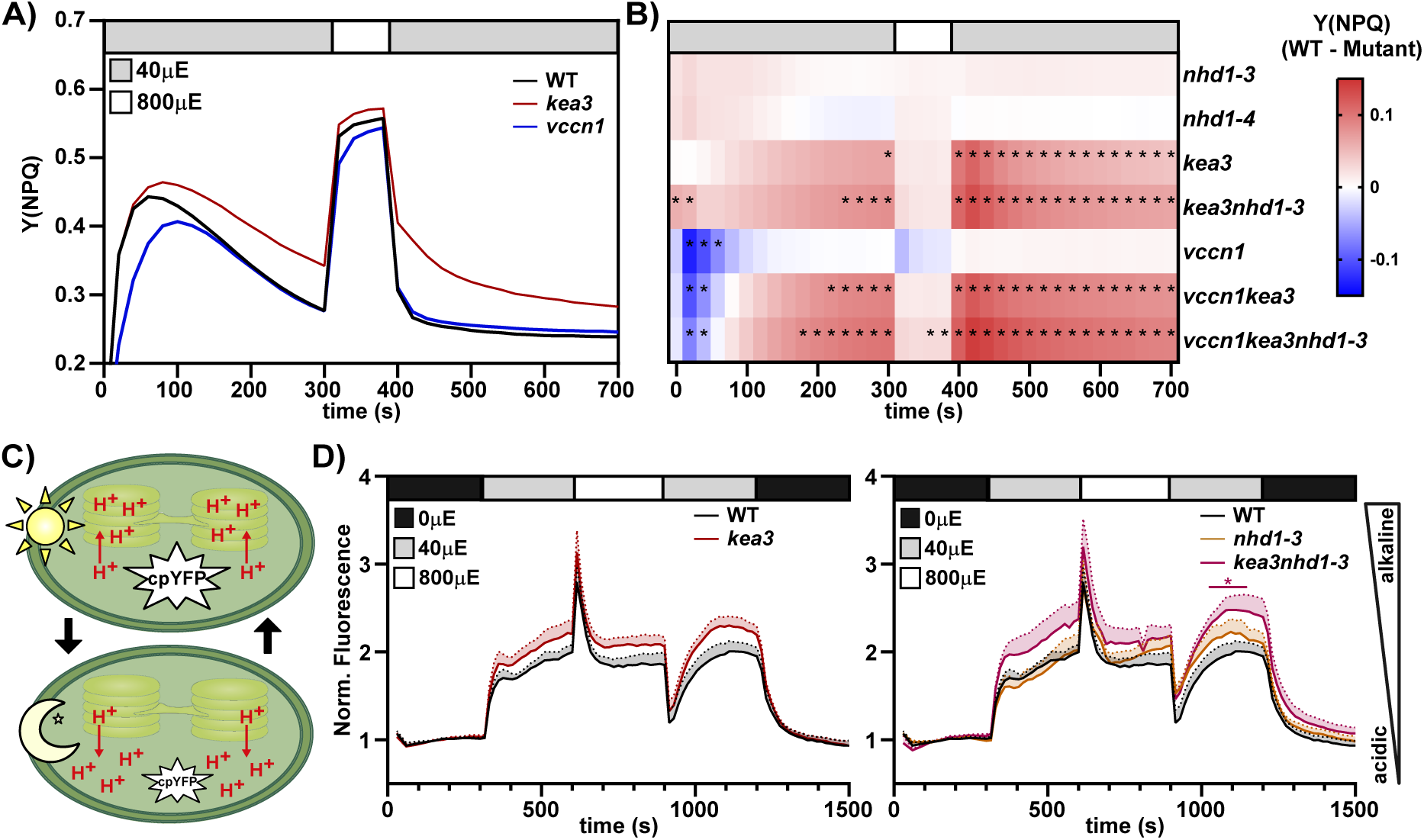
NHD1 importance is revealed in the absence of other thylakoid ion transport proteins. A) Nonphotochemical quenching (NPQ) dynamics in *A. thaliana* WT, *kea3* and *vccn1*. Plants are exposed to alternating light conditions: 300 s of low light (40 µmol photons m^−2^ s^−1^, gray box), 100 s high light (800 µmol photons m^−2^ s^−1^, white box), 300 s of low light (40 µmol photons m^−2^ s^−1^, gray box) (avg., n=8). B) Avg. changes in NPQ kinetics mutants vs. WT (shown in A) (avg., n≥9). Statistical significance was determined at each time point using a two-way ANOVA (Dunnett’s multiple comparisons test; significantly changed comparisons are indicated by an asterisk). Complete statistical analysis is provided in Table S1. C) Cartoon of fluorescence dynamics of stroma targeted circular permuted yellow fluorescent protein (cpYFP). Light treatment induces an increase in sensor fluorescence, corresponding to an increase in pH. D) Changes in stromal pH in *A. thaliana* expressing cpYFP in response to changing light conditions. Leaf discs were dark-adapted for 5 min (black box) to establish a baseline, followed by exposure to white light under alternating low light (40 µmol photons m^−2^ s^−1^, gray box), high light (800 µmol photons m^−2^ s^−1^, white box) and low light (40 µmol photons m^−2^ s^−1^, gray box) for 5 min each. Subsequently, measurements were continued for 5 min in the dark. For visual clarity the avg. ± SEM is shown (n≥4). Statistical significance was determined at each time point using a two-way ANOVA (Dunnett’s multiple comparisons test; significantly changed comparisons are indicated by an asterisk). Complete statistical analysis is provided in Table S1.

When comparing NPQ effects between genotypes, it became clear that NPQ changes from the WT were much more pronounced in the *kea3* plants than in both *nhd1* mutant lines, indicating that K^+^/H^+^ exchange across the thylakoid membrane dominates, particularly after high-to-low-light transitions (Fig. 3A, B). In contrast, *vccn1* exhibited a low Y(NPQ) induction compared to WT in the first seconds of the low light phase. This effect is thought to arise from the lack of luminal Cl^-^ influx, which delays acidification of the lumen (Davis et al., 2017).

Examining the full mutant panel showed that the effects caused by KEA3 defects are observed in all loss-of-function mutant combinations lacking this transporter. However, closer examination revealed that the NPQ response in the case of the *kea3nhd1* mutant exceeds the sum of the effects in the individual single mutants, suggesting compensatory ion flux triggered by a single loss of either KEA3 or NHD1 (Fig. 3B). A similar pattern was observed across both loss-of-function mutant combinations lacking VCCN1. Particularly during the induction phase, NPQ increased again upon loss of KEA3 in the *vccn1kea3* double mutant relative to the *vccn1* mutant. This rise in NPQ induction was further enhanced by the loss of NHD1 in the *vccn1kea3nhd1* triple mutant. Taken together, these observations suggest that the effect of NHD1 loss, albeit weak in the single mutant alleles, becomes apparent in the absence of other ion transporters.

To further elucidate the functional interplay between NHD1 and KEA3 in regulating H^+^ flux across the thylakoid, single and double mutants were transformed with the stromal pH sensor cpYFP (Zheng et al., 2025). The fluorescence intensity of cpYFP correlates with pH; for example, an increase in fluorescence indicates stromal alkalinization (Fig. 3C; Fig. S3G, H). Upon illumination, stromal cpYFP fluorescence was significantly elevated, indicating a more alkaline stromal pH (Fig. 3D). The *kea3* and *nhd1* single mutants did not show strongly altered pH responses from WT (Fig. 3D). However, both *kea3nhd1* double mutants exhibited a significant increase in stromal pH compared to the respective single mutants (Fig. 3D, right panel; Fig. S3I). Together with the photosynthetic measurements, these results suggest that loss of NHD1 affects H^+^ translocation from the thylakoid lumen to the stroma in the absence of KEA3.

### Loss of NHD1 compromises the photosynthetic response to dynamic light conditions

As a thylakoid membrane ion exchanger, NHD1 possibly plays a role in the regulation of photoprotection, an important process under dynamic light conditions as they occur in the field (Kromdijk et al., 2016; Armbruster et al., 2017; Slattery et al., 2018). Hence, *nhd1* single and higher-order loss-of-function mutants were monitored by long-term (5 days) dynamic environmental photosynthetic imaging (DEPI) under changing light conditions to reveal dynamic light-specific phenotypes (Fig. 4; Fig. S4) (Cruz et al., 2016). The *kea3* mutant showed elevated Y(NPQ), whereas the *vccn1* mutant showed reduced Y(NPQ), consistent with previous results (Höhner et al., 2019). The *nhd1-3* and *nhd1-4* plants exhibited no changes in Y(NPQ) or Y(II) as long as light intensity remained constant (days 1 and 4). In contrast, exposure to sinusoidal high-light stress (day 2) or fluctuating light treatment (days 3 and 5) resulted in a significant increase in Y(NPQ) and lower Y(II) in both *nhd1* loss-of-function alleles. Consistent with previous short-term measurements (Fig. 3B, D), the *kea3nhd1* double mutant mounted a stronger response than *kea3* single mutants. Similarly, the *vccn1kea3nhd1* triple mutant showed a further increase in Y(NPQ) compared to the *vccn1kea3* double mutant. In both comparisons, the enhanced phenotype was reflected in increased Y(NPQ) and a concomitant further reduction in Y(II). Overall, the effects of NHD1 loss on NPQ were more pronounced under dynamic light, suggesting a role for NHD1 under these conditions.

**Fig. 4.**
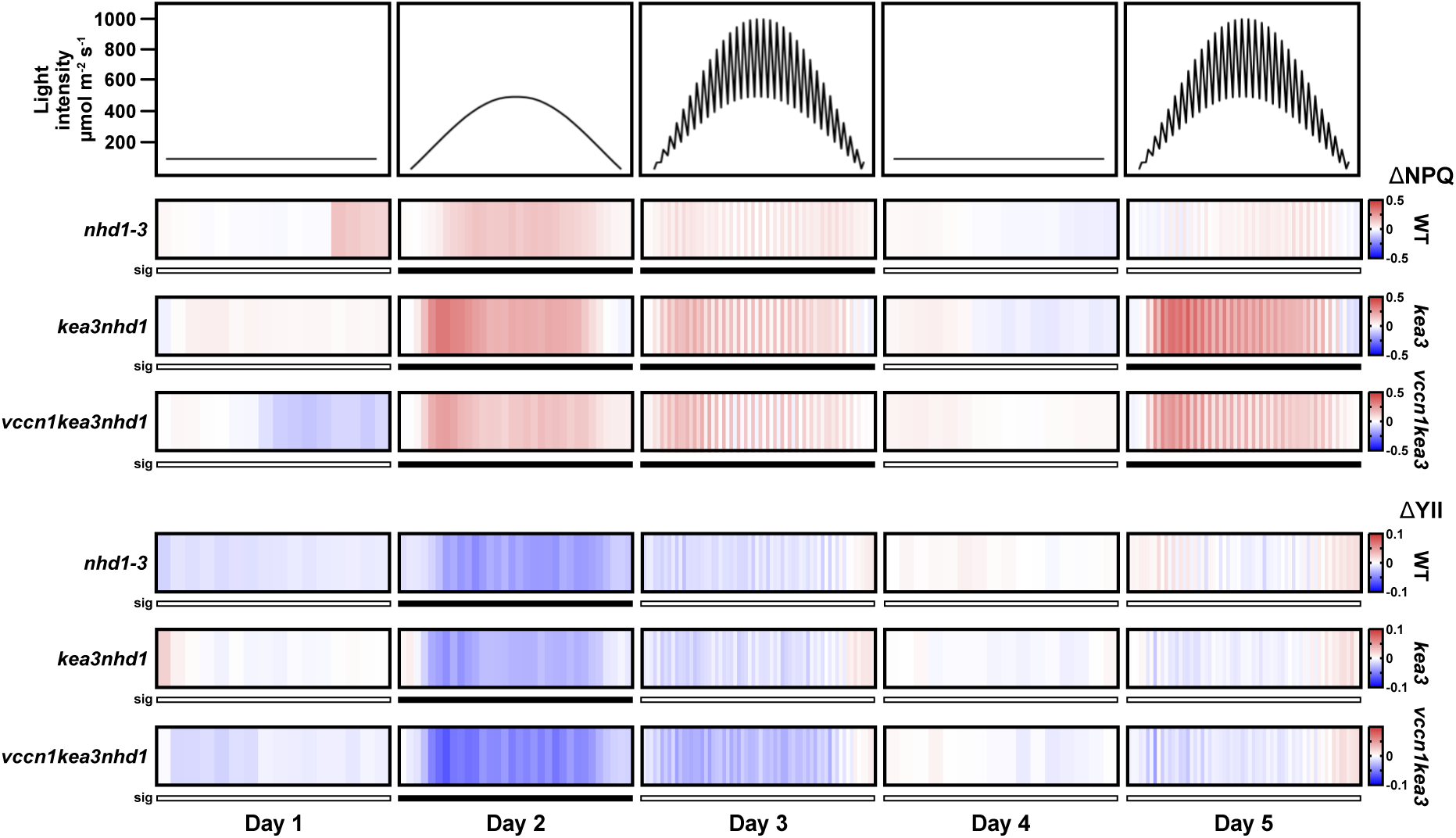
Dynamic Environment Photosynthesis Imaging (DEPI) reveals *nhd1* loss-of-function phenotypes under specific light conditions and genetic backgrounds. Plant photosynthetic performance was monitored over 5 days with different light conditions: flat day (days 1&4), sinusoidal day (day 2), fluctuating day (days 3&5). Shown are NPQ (upper half) and YII (lower half) in comparisons to WT and genetic backgrounds indicated on the right (ΔNPQ = NPQ_mutant_ - NPQ_background_) (avg., n≥10). Statistical significance was determined using a two-way ANOVA (Tukey’s post-hoc test, significantly different days are indicated by a black box). Complete statistical analysis is provided in Table S2.

### NHD1 loss-of-function mutants show a phenotype after long-term fluctuating light treatment

Encouraged by the DEPI observations, an extended long-term (14 days) fluctuating light treatment was carried out side-by-side with a standard growth light control. No genotype-specific differences were observed when plants were grown under control conditions. However, in the fluctuating light regime, *nhd1* loss-of-function mutants revealed a drastic reduction in fresh weight and rosette size (Fig. 5A). In addition to growth defects, *nhd1* mutants displayed reduced *F*_v_/*F*_m_, an indicator of photosynthetic stress and PSII impairment after fluctuating light treatment (Fig. 5A).

**Fig. 5.**
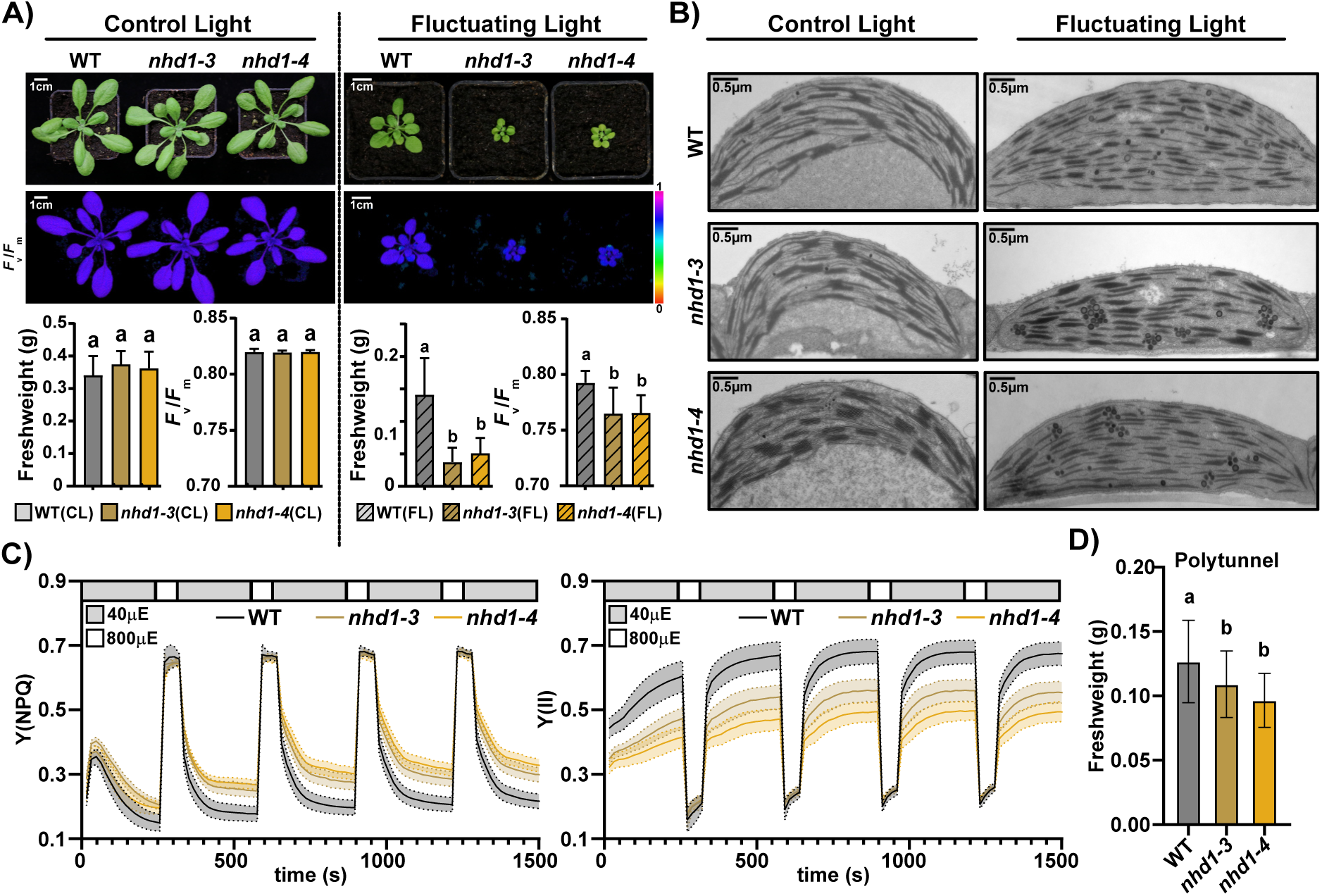
NHD1 function is crucial under long-term fluctuating light treatment. A) Photographs (upper panels) and false color images showing *F*_v_/*F*_m_ (lower panel) of 3-week-old WT, *nhd1-3* and *nhd1-4* plants after permanent growth at control and fluctuating light conditions. Below, quantification of fresh weight and the *F*_v_/*F*_m_ parameter WT and *nhd1* mutant alleles for both light conditions (avg. ± SD, n=9). Statistical significance was determined using a one-way ANOVA (*P*-value < 0.05; letters indicate significant groups). Complete statistical analysis provided in Table S3. B) Transmission-electron microscopy of WT and *nhd1* loss-of-function mutant chloroplasts. Scale bar = 0.5 µm. C) Photosynthetic performance of WT and mutant alleles after growth in fluctuating light. During the measurement, plants were exposed to alternating light conditions consisting of low light (40 µmol photons m^−2^ s^−1^, gray box, 240 s) and high light (800 µmol photons m^−2^ s^−1^, white box, 60 s) (avg. ± SD, n=9). Statistical significance was determined at each time point using a two-way ANOVA (Tukey’s post-hoc test). Complete statistical analysis see Table S3. D) Quantification of fresh weight of 3-week-old WT, *nhd1-3* and *nhd1-4* plants grown in a Polytunnel (avg. ± SD, n≥24). Statistical significance was determined using a one-way ANOVA (*P*-value < 0.05; letters indicate significant groups). Complete statistical analysis is provided in Table S3.

To investigate cellular morphological changes linked to the fluctuating light phenotype, transmission electron microscopy (TEM) was performed. Under standard growth control conditions, no obvious differences in chloroplast ultrastructure were observed across genotypes (Fig. 5B). When comparing these with plastids of fluctuating light-treated plants, it became apparent that the change in light regime affected the thylakoid ultrastructure in WT plants, increasing grana number and decreasing grana width (Fig. 5B; Fig. S5A). This is consistent with observations in field-grown *Arabidopsis* plants exposed to naturally fluctuating light conditions (Flannery et al., 2021). The two *nhd1* loss-of-function alleles showed significantly reduced grana number and width compared to the WT after fluctuating light treatment, whereas no genotype-specific difference was observed in the number of plastoglobuli (Fig. 5B; Fig. S5A).

We tracked photosynthetic parameters during fluctuating light treatments across genotypes. Both *nhd1* alleles exhibited increased Y(NPQ) and decreased Y(II) during low light, which in part explains the poor growth in these light conditions (Fig. 5C). To determine whether elevated NPQ reflects increased lumenal acidification, the q_E_ and q_I_ components were resolved. No differences in q_I_ were detected, while q_E_ was significantly increased (Fig. S5B), suggesting that the elevated NPQ in *nhd1* mutants is likely driven by enhanced proton accumulation in the thylakoid lumen. Since we were curious about the importance of NHD1 under field conditions, we also carried out an outdoor polytunnel experiment (von Bismarck et al., 2023). Although WT plants grew more slowly under natural light and temperature regimes, a well-known phenomenon (Mishra et al., 2012), both *nhd1* mutant alleles showed stronger reductions in growth (Fig. 5D).

### Mass spectrometry reveals a link between NHD1 and PSII-associated processes

Fluctuating light stress impaired photosynthetic performance and altered the thylakoid morphology in *nhd1* plants more drastically than in WT (Fig. 5; Fig. S5). To understand in what way this reshapes the plants’ proteome in the absence of NHD1, label-free quantitative (LFQ) proteomics was performed on plants before and after fluctuating light treatment (Fig. 6A). Comparison of the two light treatments revealed a gradual decline in *F*_v_/*F*_m_ over time, which was exclusive to fluctuating-light-treated plants. This decline, however, was more pronounced in both *nhd1* loss-of-function alleles, with the largest difference relative to the WT observed after four days of light treatment (Fig. 6B; Fig. S6A, B). Therefore, this time point was chosen for sample harvesting.

**Fig. 6.**
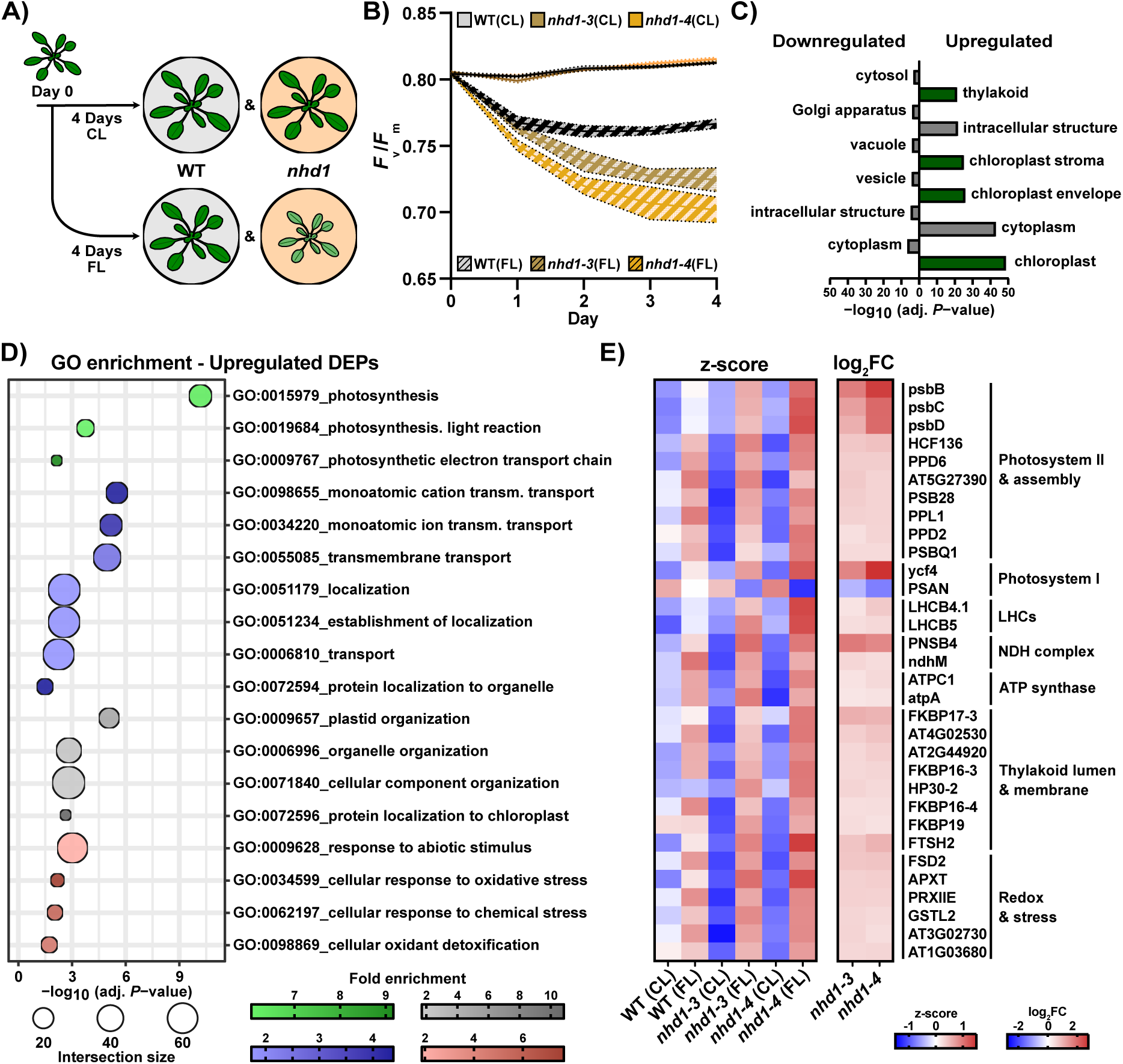
Mass spectrometry reveals reorganization of thylakoid proteins in *nhd1* mutants following fluctuating light treatment. A) Schematic overview of the experimental workflow used for mass spectrometry analysis. 2-week-old WT and *nhd1* mutants were transferred to control (CL) or fluctuating light (FL) conditions for 4 days prior to harvest. Proteomic changes were analyzed within each genotype to characterize the acclimation response to fluctuating light. B) *F*_v_/*F*_m_ measurements during light treatments show most dramatic PSII impairment after 4 days i.e., timepoint of sampling. C) Localization of differentially expressed proteins (DEPs) (adj. *P*-value < 0.05). D) Selected significantly enriched GO-terms (Benjamini-Hochberg FDR < 0.05). Circle size corresponds to the number of DEPs within GO-terms, while coloring depicts fold enrichment of manually selected groups. More DEPs included in GO-terms are provided in Table S4. E) Heatmap depicts z-scores of lable free protein quantification values (left) and log_2_FC values of mutant lines vs. WT (right) of selected thylakoid-related proteins.

Analysis of the acclimation response to fluctuating light identified 2,641 differentially expressed proteins (DEPs) irrespective of the genotype, highlighting the substantial impact of the light regime itself on the proteome. Of these, 592 were exclusively found to be changed in both *nhd1* mutants (Fig. S6C). These included 412 upregulated and 180 downregulated proteins. According to SUBA Multiple Marker Abundance Profiling (MMAP) (Hooper et al., 2017), the majority of upregulated *nhd1-exclusive* DEPs reside in the chloroplast and its subfractions, while downregulated proteins were scattered across other compartments (Fig. 6C). As we intended to focus primarily on changes in the chloroplast, the analysis from this point forward focused on upregulated proteins. Biological functions of upregulated *nhd1*-exclusive DEPs were inferred through Gene Ontology (GO) enrichment analysis (Fig. 6D, Fig. S6D). A first set of enriched GO terms was related to photosynthesis. These terms included “photosynthesis”, “photosynthesis, light reaction,” and “photosynthetic electron transport chain”. Since NHD1 is located in the thylakoid membrane (Fig. 1), an altered response of photosynthesis-associated proteins relative to WT appears plausible when NHD1 is missing. In addition to this group, GO terms such as “monoatomic cation transmembrane transport”, “plastid organization” and “response to abiotic stress” are also significantly enriched, hinting at wide-ranging changes throughout the plant cell and the chloroplast of *nhd1* mutants. Upon closer examination of thylakoid-localized proteins that were significantly upregulated only in *nhd1* mutants, several PSII-associated proteins can be detected (Fig. 6E), suggesting that lack of NHD1 results in exacerbated PSII-related mechanisms. In addition, differences in the abundance of proteins involved in cyclic electron flow (CEF) and ATP synthase were observed in the *nhd1* mutants, potentially indicating compensatory adjustments for proton flux.

### Simulating NHD1 Na^+^/H^+^ antiport using a model of photosynthetic light reactions

PMF models enable the quantitative assessment of proton and ion fluxes across the thylakoid membrane, however, this requires incorporating all ion transport processes contributing to PMF formation, partition and regulation (Davis et al., 2017). In this work, we have shown that NHD1 is a thylakoid-localized Na^+^/H^+^ antiporter that contributes to photosynthetic acclimation under long-term fluctuating light conditions. Hence, we expanded a computational model of photosynthesis to a third ion species, Na^+^ (Fig. 7A). A detailed description of the model is provided in Ebeling et al. (2026).

**Fig. 7.**
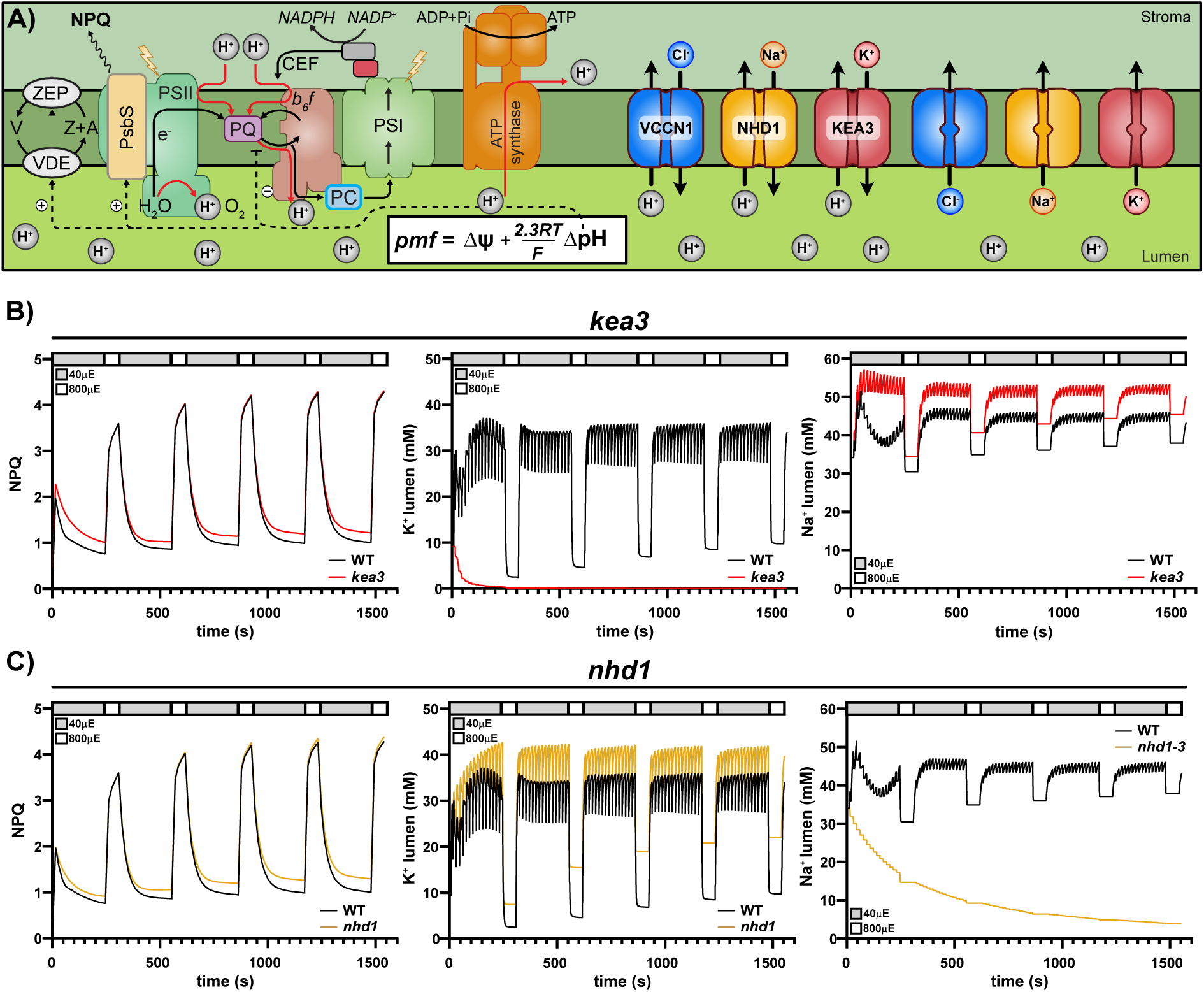
Photosynthesis simulation in the presence or absence of thylakoid K^+^/H^+^ and Na^+^/H^+^ antiport, respectively. A) Extended photosynthesis model based on Ebeling et al., 2026, with Na^+^ as a third metal ion. Simulation of WT vs. B) *kea3* and C) *nhd1* loss-of-function mutant phenotypes under fluctuating light conditions. Modelled parameters: NPQ (left), lumen K^+^ (middle) and lumen Na^+^ concentration (right).

In the model, we described NHD1 as an antiporter that imports Na^+^ into the lumen in exchange for protons. NHD1 is proposed to be active during transitions from high to low light. Thus, to model this activity, we used the redox status of the PQ pool as an approximation for (de-)activation. We also included a cation channel to ensure that Na^+^ can return to the stroma. This hypothetical cation channel is modeled similarly to VCCN1, opening and closing at a certain voltage to allow sodium to diffuse back into the stroma. To assess model performance under long-term dynamic light conditions, we simulated repeated transitions between low and high light (40 µE 4 min and 800 µE 1 min) for three scenarios: WT, *kea3* and *nhd1* (Fig. 5).

The model accurately reproduced the experimentally measured NPQ dynamics of WT plants (Figs. 5C and 7B, C). Moreover, *in silico* removal of the K^+^/H^+^ antiporter recapitulated the previously reported *kea3* NPQ phenotype (Fig. 7B). In the *kea3* model, the increase in NPQ was accompanied by depletion of luminal K^+^ and a concomitant accumulation of luminal Na^+^, suggesting that Na^+^ flux by NHD1 may provide a compensatory ionic flux in the absence of K^+^ transport. The experimentally observed *nhd1* loss-of-function phenotype was simulated by omitting the sodium antiporter from the model. Notably, the simulated phenotype matches the measured NPQ kinetics of *nhd1* mutant alleles (Figs. 5C and 7C). According to the simulation, an absence of NHD1 leads to a continuous decline in the luminal sodium concentration, while NPQ increases (Fig. 7C). Consequently, the phenotype arises from a decreased capacity to export protons into the stroma in a timely fashion (Fig. S7A, B). This deficit results in lower luminal pH and, therefore, more persistent NPQ. Furthermore, the simulations indicate that the loss of Na^+^/H^+^ antiporter activity affects the luminal concentrations of other ions as well, as the K^+^ concentration in the lumen increases throughout the measurement (Fig. 7C). These results suggest the presence of compensatory transport mechanisms that help to buffer the loss of antiporters, while further highlighting the complex nature of PMF regulation.

## Discussion

The current conceptual framework around thylakoid ion homeostasis has so far been focused on protons, K^+^, and Cl^-^ and their relation to photosynthetic control and lumen acidification. In this study, we have demonstrated that NHD1 exchanges Na^+^ for protons across membranes and resides in the thylakoid. Indeed, this work showed not only that Na^+^ is present in physiological amounts in the chloroplast, but also that the loss of NHD1 severely affects plant performance under natural dynamic light conditions. Elevated NPQ, PSII impairment, and significantly reduced growth rates of *nhd1* loss-of-function mutants support this claim, indicating an imbalance between light absorption and downstream energy utilization.

It is noteworthy that, except for altered ECS inversion kinetics, *nhd1* plants show virtually no differences from WT when grown under control light. After fluctuating light treatment or in higher-order mutant backgrounds, however, *nhd1* loss-of-function mutants show delayed NPQ relaxation as well as reduced Y(II) akin to *kea3* plants. Also, kinetic modeling of both antiporter null alleles revealed similar, yet not identical, NPQ responses. Despite NHD1 and KEA3 both being cation/H^+^ antiporters, one should not necessarily expect them to be functionally redundant. The crucial question is: Why should Na^+^/H^+^ exchange only matter after prolonged fluctuating light exposure? There are three non-mutually exclusive possibilities: 1) NHD1 exchange activity may be lower than KEA3 activity under “normal” conditions *in planta*. Even though we measured *in vitro* antiport, H^+^ flux also depends substrate concentrations, on protein abundance, and the activation state of the transporter. 2) NHD1 may be regulated differently from KEA3. To contribute to acute NPQ relaxation mechanisms, tight light-responsive regulation is needed. The KEA3 C-terminus was described to respond to the energy state of the chloroplast stroma, thereby regulating transport activity in a redox-dependent manner (Uflewski et al., 2024). Measurements of NHD1’s pH dependence revealed a loss of activity under alkaline conditions. This is supported by previous work on Na^+^/H^+^ antiporters, in which charged and conserved residues at the substrate tunnel entrance of NhaD transporters were described as a “pH sensor”, rendering transport activity pH-sensitive (Lv et al., 2022). If the pH-sensitive site of the transporter is oriented toward the stroma, this would result in transporter inactivation upon stromal alkalization. Also, the involvement of additional regulatory sites cannot be excluded. 3) NHD1-driven Na^+^/H^+^ exchange may be more important when repeated light fluctuations, as they occur outside, alter the ion homeostasis in chloroplasts. However, under control light NHD1 seems to contribute little to transthylakoid ion flux and other fast-acting mechanisms may compensate for its absence. This is supported by the spectroscopic alterations observed in the *kea3nhd1* double mutants, which were stronger than changes in the individual mutants in both NPQ and pH measurements. Growth under permanent fluctuating light, however, could alter overall ion homeostasis, rendering NHD1 activity necessary for the photosynthetic acclimation response and explaining the diverse mutant phenotypes that emerge from a loss of NHD1. Hence, Na^+^/H^+^ antiport across the thylakoid membrane of plants seems to represent an adaptive buffering system to natural light conditions, explaining why its function has been retained.

Since no NHD1 homologue was found in Cyanobacteria so far, it is puzzling how the genes encoding these transporters emerged in so many Chloroplastida genomes (Kunz et al., 2024). Interestingly, the NhaD-type carrier NHD1 is among the plastid proteins encoded by Chlamydiae-derived genes (Moustafa et al., 2008). Chlamydiae, the obligate intracellular bacteria of protists, have been debated for decades as contributors to establishing endosymbiosis (Domman et al., 2015; Cenci et al., 2017; Maire et al., 2024). This may suggest that transthylakoid Na^+^/H^+^ exchange was not lost during chloroplast evolution but was instead functionally conserved through a substitution of the cyanobacterial NhaS3 linage by a Chlamydiae-derived NhaD transporter. In this scenario, the functional redundancy created by the addition of another Na^+^/H^+^ antiporter may have supported the functional divergence of transthylakoid ion transport toward K^+^/H^+^ exchange, now performed by KEA3, the closest homologue of NhaS3 (Tsujii et al., 2019). This would imply strong selective pressure to retain Na^+^/H^+^ flux across the thylakoid membrane.

Importantly, if NHD1 homologues localize exclusively to the thylakoid membrane across C_3_ and C_4_ monocots and dicots, other transport mechanisms must exist in the envelope to balance Na^+^ influx into plastids via BILE ACID:SODIUM SYMPORTER (BASS) proteins, which import pyruvate, glycolate, and glucosinolate (Gigolashvili et al., 2009; Furumoto et al., 2011; South et al., 2017).

In summary, this work positions NHD1-mediated Na^+^/H^+^ exchange across the plant thylakoid membrane as a novel “conserved layer” of luminal ion homeostasis localizing alongside known transporters and channels of the thylakoid membrane (Fig. 8). Comparing NHD1 function to that of KEA3, the following mechanistic separation could be made: KEA3 regulates the major rapid response to changes in light intensity, whereas NHD1 contributes to long-term acclimation of thylakoid ion homeostasis to light fluctuations. Our work suggests that Na^+^ itself has a more fundamental role in plant thylakoid physiology than previously assumed.

**Fig. 8.**
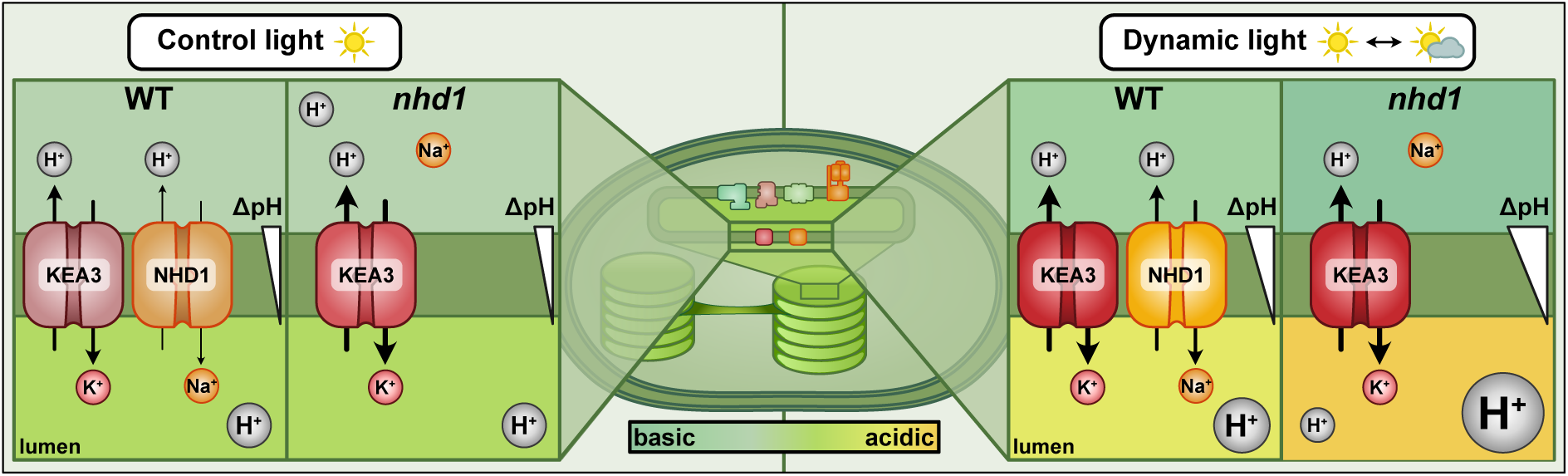
Working model: The thylakoid membrane Na^+^/H^+^ antiporter NHD1 and its role in acclimation to dynamic light stress. Na^+^/H^+^ exchange of the thylakoid membrane provides an additional export route for protons particularly important under extreme light conditions (e.g., prolonged changes in light intensity). Fluctuating light treatment and *nhd1* loss-of-function results in a higher ΔpH, causing increased NPQ and a decrease in YII compared to the WT. Na^+^/H^+^ exchange across the thylakoid links suborganellar Na^+^ homeostasis and long-term acclimation of photosynthesis.

## Materials and Methods

### Plant growth

*Arabidopsis thaliana* accession Columbia-0 (Col-0; WT) and mutant plants were germinated and grown on soil (Substrate yellow line, Stender, Schermbeck, Germany) under long-day conditions (16 h light/8 h dark). Plants were maintained at 22°C and 50% humidity under a light intensity of 100 µmol photons m^−2^ s^−1^. Unless otherwise indicated, rosettes from 3-week-old plants were used for all experiments.

For fluctuating light experiments, plants were germinated and grown for one week at 100 µmol photons m^−2^ s^−1^ under long-day conditions (16 h light/8 h dark). One group was maintained under constant illumination at 100 µmol photons m^−2^ s^−1^, whereas the second group was exposed to fluctuating light conditions in a custom-made illumination system (Schneider et al., 2019). The fluctuating light regime consisted of alternating 1 min periods of high light (800 µmol photons m^−2^ s^−1^) and 4 min periods of low light (40 µmol photons m^−2^ s^−1^) under long-day conditions (16 h light/8 h dark).

### Isolation of single and higher-order mutants

Higher-order mutants were obtained through crosses between the following mutant lines: *nhd1-3* (kindly provided by Czaba Koncz (Koncz et al., 1992)), *nhd1-4* (SALK_118891), *kea3-1* (SAIL_556_E12 (Kunz et al., 2014)), *kea3-1* (Gabi_170G09 (Armbruster et al., 2014)) and *vccn1-1* (SALK_103612C). Double mutants were generated by crossing *kea3-1* (SAIL_556_E12) with *nhd1-3* and *kea3-1* (Gabi_170G09) with *nhd1-4*. The triple mutant *kea3-1vccn1nhd1* was obtained by crossing *kea3-1vccn1* with *kea3-1nhd1-3*. Homozygous T-DNA lines were confirmed by PCR (Figs. S1A, B and S3C–E).

### Confocal microscopy for protein localization studies

Images were acquired using a Leica Stellaris 5 Confocal Laser Scanning Microscope (Leica, Wetzlar), equipped with a 405 nm diode and a supercontinuum White Light Laser (WLL). Emission was detected with a PowerHybridHyd S detector. For protein localization studies, yellow fluorescent protein (YFP) was excited at a wavelength of 515 nm and red fluorescent protein (mCherry) was excited at a wavelength of 587 nm, while chlorophyll autofluorescence was excited at 405 nm. Emission was detected at 520–580 nm for YFP, 596–622 nm for mCherry, and 623–813 nm for chlorophyll. A z-stack of 13 images was acquired and processed using the LAS X software to create the maximum projection.

### Chloroplast isolation for further analyses

Intact chloroplasts were isolated according to Aronsson and Jarvis (2002). Briefly, 3-week-old *Arabidopsis* plants were dark-adapted for 24 h prior to harvest. Plant material was homogenized in pre-cooled isolation buffer (300 mM sorbitol, 5 mM MgCl_2_, 5 mM EDTA, 20 mM HEPES/NaOH, pH 8.0) using a Polytron homogenizer (Bachhofer, Reutlingen, Germany), filtered through gauze, and centrifuged at 1500*g* for 5 min at 4°C. The resulting pellet was resuspended in isolation buffer and layered onto a two-step Percoll gradient consisting of 82% (v/v) and 30% (v/v) Percoll (GE Healthcare, Chicago, USA) prepared in the same buffer. After centrifugation at 3500*g* for 10 min at 4°C, intact chloroplasts were collected from the interface, washed twice in isolation buffer, and pelleted at 1500*g* for 5 min at 4°C.

Chloroplasts for quantification of plastid Na^+^ levels were isolated in a salt free isolation buffer (300 mM sorbitol, 5 mM Mg(CH_3_COO)_2_, 5 mM EDTA, 20 mM MOPS/Tris, pH 8.0). Dried aliquots of chloroplast suspensions and isolation buffer were digested in 2 mL 65% (w/w) HNO_3_ and 1 mL 30% (w/w) H_2_O_2_ (both analytical grade) inside acid-washed 55-mL PFA vessels of a Mars5 Xpress microwave digestion system (CEM, Kamp-Lintford, Germany), by keeping at RT for 30 min, heating to 190°C within 45 min, holding at 190°C for 15 min, followed by passive cooling to RT. Samples were transferred into 15-mL round-bottom screw-cap polypropylene tubes and filled up to a total volume of 10 mL with ultrapure water. Sodium concentrations were quantified at a wavelength of 589.592 nm (radial) using Inductively Coupled Plasma-Optical Emission Spectrometry (ICP-OES, Agilent 5800VDV, Agilent Technologies, Waldbronn, Germany), with baseline correction according to isolation buffer samples. Calibration was achieved using a concentration series of multi-element standards, and measurement accuracy was verified through intermittent measurements of blanks and of one multi-element calibration standard solution.

For sub-chloroplast fractionation (Bouchnak et al., 2018), isolated chloroplasts were disrupted in hypotonic burst buffer (5 mM MgCl_2_, 20 mM HEPES/NaOH, pH 8.0) and layered on a three-step sucrose gradient (10 mL of 0.46 M, 10 mL of 1.00 M, and 6 mL of 1.20 M sucrose). After centrifugation for 2 h at 58,000*g* in a swing-out rotor, the stroma, envelope and thylakoid fractions were collected, washed several times, resuspended in burst buffer and stored until further usage in immunoblots.

### Protein extraction and immunoblotting

Plant material was normalized based on fresh weight, homogenized and solubilized for 20 min at 4°C in solubilization buffer (10% (v/v) glycerol, 150 mM NaCl, 2.5 mM EDTA, 10 mM DTT, 50 mM Tris-HCl pH 7.5, 1.5% (w/v) SDS, 1% NP40 (v/v), plant protease inhibitor mix). Samples were then centrifuged at 10,000*g* for 10 min at 4°C, mixed with SDS loading dye (100 mM DTT, 62.5 mM Tris-HCl pH 6.8, 2% (w/v) SDS, 10% (v/v) glycerol, 0.003% (w/v) bromophenol blue), and heated for 15 min at 70°C. Proteins were separated by SDS-PAGE (10% acrylamide) and transferred to PVDF membranes in a wet-blotting system. Membranes were incubated overnight at 4°C with the respective primary antibodies: α-AtNHD1 (Lopez et al., 2022), α-PsaF (AS06104), α-PsbS (AS03032) and α-KEA1/2 (Bölter et al., 2020). The blot was rinsed three times for 10 min each with TBST and subsequently incubated with horseradish peroxidase (HRP)-conjugated secondary antibody. Enhanced chemiluminescence (ECL) was used for detection on an ImageQuant LAS 4000 system (GE Healthcare, Chicago, USA).

### Plasmid construction

The NhaS3, KEA3, and NHD1 inserts were amplified from cDNA using the primers listed in Table S1. Restriction-ligation was performed by digesting both the pACBAA_His vector, a modified pACBAD14 vector in which chloramphenicol was swapped for ampicillin resistance (Wunder et al., 2026), and the PCR-amplified inserts with NcoI and XhoI for NhaS3 and NHD1 insertion or with NcoI and NotI for KEA3 insertion, followed by ligation using T4 DNA ligase (NEB). The resulting plasmids were designated pACBAA_NhaS3+His, pACBAA_KEA3+His and pACBAA_NHD1+His.

### Complementation assays and antiporter assays

For complementation and antiporter assays, the *E. coli* strains TO114 (W3110 *nhaA*::Km^r^ *nhaB*::Em^r^ *chaA*::Cm^r^) (Ohyama et al., 1994) and LB2003 (F-, *thi*, *lacz, gal*, *rha*, Δ*kdpFABC5*, *trkD1*, Δ*trkA*) (Stumpe and Bakker, 1997) were transformed with the empty vector or the respective plasmids. *E. coli* cells were cultured in liquid medium starting at an OD of 0.1. Gene expression was induced 30 min later by the addition of 0.2% arabinose, and cells were harvested after two additional hours. After incubation, cells were collected by centrifugation, washed three times with water, and plated as follows. To investigate Na^+^ transport, *E. coli* TO114 transformants were grown on LBK (1% (m/v) peptone, 0.5% (m/v) yeast extract and 100 mM KCl) with defined NaCl concentrations (0 or 300 mM). To investigate K^+^ transport, *E. coli* LB2003 transformants were grown on LB (1% (m/v) peptone, 0.5% (m/v) yeast extract) with defined KCl concentrations (0 or 100 mM).

For the isolation of inverted vesicles (Rosen and Tsuchiya, 1979), DH5α (WT) and *E. coli* transformants were grown in LBK medium at 37°C until an OD of 0.6 was reached. Arabinose was then added to a final concentration of 0.2% to induce expression, and cultures were incubated for an additional 2 h. Cells were harvested at 5000*g* for 10 min, washed twice with press medium (10 mM MOPS-Tris buffer pH 7.5, 300 mM choline chloride, 25 mM MgSO_4_), and then resuspended in the same buffer supplemented with 0.5 mM phenylmethylsulfonyl fluoride and 20% glycerol. The bacterial suspension was then passed through a French press at 1000 psi twice. Unbroken cells were removed by centrifugation at 10,000*g* for 10 min. The membrane fraction was collected by ultracentrifugation at 100,000*g* for 60 min at 4°C and then resuspended in fresh press buffer. The protein concentration was determined using the Bradford assay with bovine serum albumin (BSA) as a standard. Membrane preparations were stored at −80°C.

Cation antiporter activity was determined by measuring acridine orange fluorescence dequenching (Holdsworth and Law, 2014). The assay mixture contained 200 µL of 10 mM Tris-HCl (pH 7.0-9.0), 140 mM choline chloride, 5 mM MgSO_4_, 1 µM acridine orange, and 5 µg of vesicle protein. Respiration-dependent formation of the transmembrane pH gradient was initiated by the addition of 10 mM Tris/D-lactate. The resulting quenching of acridine orange fluorescence was monitored using a TECAN Spark plate reader (Tecan, Männedorf, Switzerland) at an excitation wavelength of 492 nm and an emission wavelength of 525 nm. Cation-dependent H^+^ efflux from the vesicle lumen was monitored as fluorescence dequenching of acridine orange. At the end of the assay, 25 mM NH_4_Cl was added to dissipate the remaining proton motive force (PMF). NaCl, KCl, LiCl, and RbCl were used at 10 mM unless stated otherwise. The percentage of dequenching was calculated relative to the total respiration-dependent fluorescence quenching.

### Photosynthetic measurements

All experiments were performed at 22°C using 3-week-old soil-grown plants that were dark-adapted for 30 min prior to measurement. Chlorophyll fluorescence parameters were measured using an Imaging-PAM system (Walz GmbH, Effeltrich, Germany). Standard induction curves were carried out at a light intensity of 110 µmol photons m^−2^ s^−1^. PSII properties under short-term fluctuating light were determined under the light intensities described. Energy-dependent quenching (q_E_), and photoinhibitory quenching (q_I_) were estimated as described previously (Kramer and Crofts, 1996; Baker et al., 2007). The extent of the q_I_ component of NPQ was determined following 15 min of dark relaxation.

### ECS

Electrochromic shift (ECS) measurements were performed on a custom-built spectrophotometer (Hall et al., 2013) as in Davis et al. (2016). The ECS measurements were normalized to total pigment content extracted from equally sized leaf discs in acetone to quantify chlorophyll (Porra et al., 1989).

### pH sensor measurements

Dynamics during light transition were monitored using a Zeiss LSM 980 confocal laser scanning microscope (Carl Zeiss Microscopy GmbH, Jena, Germany) equipped with a 10x lens (Plan-Apochromat, 0.3 N.A.). A customized lightbox (Zheng et al., 2025) was manually switched between two light intensities (high light: 800 and low light: 40 µmol photons m^−2^ s^−1^). Light was provided by a cold-white LED (Max Pferdekaemper GmbH & Co. KG, Menden, Germany) implemented in the lightbox (Zheng et al., 2025). Confocal imaging was performed using discs excised from true leaves of 4-to 5-week-old *Arabidopsis thaliana* plants. The leaf discs were mounted in perfluorodecalin between two coverslips and the LED was directly attached from the top side. For pH measurements, cpYFP fluorescence was excited at 488 nm, while emission was collected at 508–535 nm. Chlorophyll fluorescence was collected simultaneously at 650–700 nm. Resolution was set to 512 × 512 pixels and 0.6475 s frame time, and pinhole to 142 µm. Plants were dark-acclimated for at least 30 min before starting the measurements. Fluorescence levels were collected from the entire microscopy frame and baseline-corrected using the dark phase (before the light treatment). To ensure that pH changes were measured in photosynthetically active plastids, the focal plane was placed in the mesophyll chloroplasts of the subepidermal adaxial leaf side, the palisade parenchyma (Zheng et al., 2025).

### Dynamic Environmental Photosynthetic Imaging (DEPI)

Homozygous loss-of-function lines were monitored for a period of five consecutive days using dynamic light intensities as described earlier (Cruz et al., 2016).

### Transmission electron microscopy (TEM)

Fixation followed the same procedure as described previously (Brandt et al., 2026). Leaves were cut into 1 mm^2^ pieces in fixation buffer (75 mM cacodylate, 1mM MgCl_2_, pH 7.0) supplemented with 2.5% glutaraldehyde and incubated for several days until further processing. Following postfixation with 1% OsO_4_, the samples were contrasted en bloc with 1% uranyl acetate in 20% acetone and dehydrated with a graded acetone series. After infiltration with Spurr’s resin, the samples were embedded and polymerized at 63°C.

### Mass spectrometry

Sample preparation was performed as described previously (Brandt et al., 2026). Briefly, for each biological replicate (four per genotype and condition), 100 mg of leaf tissue was snap-frozen in liquid nitrogen and ground to a fine powder. Proteins were extracted in 300 µL of extraction buffer (100 mM HEPES, pH 7.5, 150 mM NaCl, 10 mM DTT, 6 M guanidine hydrochloride, 1× Complete™ protease inhibitor cocktail; Roche), followed by sonication (three cycles of 20 s; Branson Sonifier B-12) and incubation at 60°C for 10 min. Following centrifugation at 10,000*g* for 15 min, proteins were precipitated using methanol/chloroform/water (4:2:3, v/v/v). The protein-containing interphase was collected, washed five times with methanol, air-dried, and resuspended in 50 mM HEPES (pH 7.8) containing 3 M urea and 2 M thiourea. Protein concentration was determined using the Pierce 660 nm Protein Assay (Thermo Fisher Scientific). For digestion, 100 µg of protein was reduced with 10 mM DTT for 30 min at 37°C, alkylated with 50 mM iodoacetamide for 20 min at room temperature in the dark, and digested with 1 µg of sequencing-grade trypsin (Thermo Fisher Scientific) overnight at 37°C. Peptides were desalted using C18 StageTips (Rappsilber et al., 2003).

Liquid chromatography–tandem mass spectrometry (LC–MS/MS) was performed using a nano-LC system (nanoElute 2, Bruker Daltonics) coupled to a timsTOF HT mass spectrometer (Bruker Daltonics) equipped with a CaptiveSpray nano-electrospray ionization (ESI) source. The nano-LC system was fitted with a PepSep Ultra separation column (C18, 25 cm × 75 µm, 1.5 µm particle size; Bruker Daltonics). Peptides (200 ng) were separated at a constant flow rate of 300 nL/min using the following gradient of mobile phase B: 2–3% B (0–1 min), 3–17% B (1–36 min), 17–25% B (36–60 min), and 25–37% B (60–66 min), followed by an increase to 95% B (66–67 min) and a 10-min wash at 95% B (67–77 min). The column temperature was maintained at 50°C throughout the analysis. Data-independent acquisition (DIA) was performed using the following settings: mass range, 100–1700 m/z; ion mobility range, 0.85–1.30 V·s/cm²; ramp time, 100 ms.

Mass-spectrometry raw data were deposited to the ProteomeXchange Consortium via the PRIDE partner repository (Perez-Riverol et al., 2022) with the identifier PXD087607. Raw data were processed using DIA-NN v2.0 (Demichev et al., 2020). A library-free analysis was performed using the *Arabidopsis thaliana* reference proteome (UniProtKB UP000006548; March 2025) supplemented with a common-contaminants database (cRAP). An in silico-predicted spectral library was generated, followed by a first-pass analysis of the DIA data to build an experiment-specific spectral library and a second-pass reanalysis of all 24 runs. Trypsin specificity was applied, allowing up to one missed cleavage. Carbamidomethylation of cysteine residues was set as a fixed modification. Match-between-runs was enabled. MS1 and MS2 mass accuracies were inferred automatically by DIA-NN. False discovery rates (FDRs) were controlled at 1% at both the precursor and protein-group levels using the DIA-NN target–decoy approach. Further analysis was conducted in Perseus (Tyanova et al., 2016). Only protein groups with valid quantitative intensity values in at least three of four biological replicates in at least one condition were retained. Missing values were imputed using values drawn from a normal distribution. Differentially expressed proteins were determined in the Perseus software with standard settings.

### GO term enrichment

To assess enriched Gene Ontology (GO) terms, gene identifiers corresponding to significantly different protein abundances (adjusted *P*-value <0.05) in *nhd1-3* and *nhd1-4* when compared to WT were pasted into the web portal of g:Profiler (https://biit.cs.ut.ee/gprofiler) (Kolberg et al., 2023) using *Arabidopsis thaliana* with default values. To condense significantly enriched GO terms (see Table S5), these terms and their *P*-values were submitted to the REVIGO web server (http://revigo.irb.hr/) (Supek et al., 2011). The resulting network was exported, imported into Cytoscape (v3.9.1) (Shannon et al., 2003), and nodes were recolored and resized according to the REVIGO *P*-values and sizes of GO terms, respectively. Bubble plots were created using ggplot2 (Wickham, 2016) in the R environment.

### Mathematical modeling of NHD1-mediated ion transporter

The model of photosynthesis utilized here is based on Ebeling et al., 2026. The full model description is in the corresponding publication. We applied several changes between the published version and the one used here. To this model version, we added sodium as an additional ion species. To regulate the sodium ions, we include a voltage-gated channel that works similarly to VCCN1 but allows Na⁺ flux into the stroma. NHD1 is assumed to have a similar activation mechanism to KEA3, but instead of using ATP and stromal pH for activation, we use Plastoquinone as a redox-status proxy of the system; when the system relaxes, NHD1 activates. The activation of both the antiporter and the voltage-gated sodium channel follows the same mathematical concept of a sigmoid activation function. Additionally, we used different numerical values for the stoichiometries of ions transferred between the stroma and lumen, without incorporating stroma and lumen volume. Both versions, including volume or not, can replicate the NPQ measurements equally well.

### Statistical analysis and figure design

Data were analyzed using GraphPad Prism (Dotmatics, Boston, USA). Significance was determined by one-way ANOVA or two-way ANOVA followed by Tukey’s multiple-comparison test unless stated otherwise.

Figures were built using Affinity Designer (Serif, Nottingham, UK).

### AI use disclosure

The authors used Anthropic’s Claude Sonnet 4.6 to assist with language editing and phrasing during the revision of the manuscript. All AI-assisted outputs, including derived output tables, were critically checked and verified by the authors against the underlying data. AI was not used to generate experimental data, and all scientific interpretation and conclusions remain the responsibility of the authors.

## Accession numbers

*NHD1* (At3g19490), *KEA3* (At4g04850), *VCCN1* (At3g61320), *nhd1-3* (Czaba Koncz) (Koncz et al., 1992), *nhd1-4* (SALK_118891), *kea3-1* (SAIL_556_E12), *kea3-1* (Gabi_170G09), *vccn1-1* (SALK_103612C).

## Acknowledgements

We thank Drs. Begoña Benito Casado (Universidad Politécnica de Madrid) and Nobuyuki Uozumi (Tohoku University) for providing *E. coli* mutant strains and Dr. Klaas von Wijk (Cornell University) for early consultation on NHD1 proteomics data in thylakoid fractions. Furthermore, we acknowledge Drs. Ricarda Höhner, Chantal Priesnitz and Carsten Völkner (At WSU previously), and Ms. Dunia-Carina Velazquez-Ayala (now at MPIIB Berlin) for assistance with the first NHD1 antibody genotyping, and polytunnel experiments, respectively. Dr. Koncz from MPI Cologne kindly provided the *nhd1-3* allele from his own T-DNA collection.

## Author Contributions

H.-H.K. conceived the research idea. H.-H.K. and E.W. wrote the manuscript. E.W. carried out the majority of experiments, analyzed data, and designed figures. B.S. assisted with experimental work. S.M. performed confocal laser microscopy and assisted with figures. B.Bö. and B.Br. fractionated and immunoblotted plastid membranes. G.A.D. cloned the NhaS3 construct. G.A.D. and D.D.S. performed ECS and analyzed data. L.H. and U.K. contributed element analyses. P.B. and M.S. assayed *in vivo* pH shifts. Q.H.N. and U.A. ran DEPI and analyzed data. C.S. and A.K. took TEM images. S.S. performed MS and analyzed data. H.J. supervised isolation of inverted vesicles. J.E. and A.M. modeled light-dependent behavior of mutants. All authors edited the manuscript.

