## Supplemental Figures for "Na^+^/H^+^ antiport across the thylakoid membrane is functionally conserved in the Chloroplastida *Arabidopsis*"

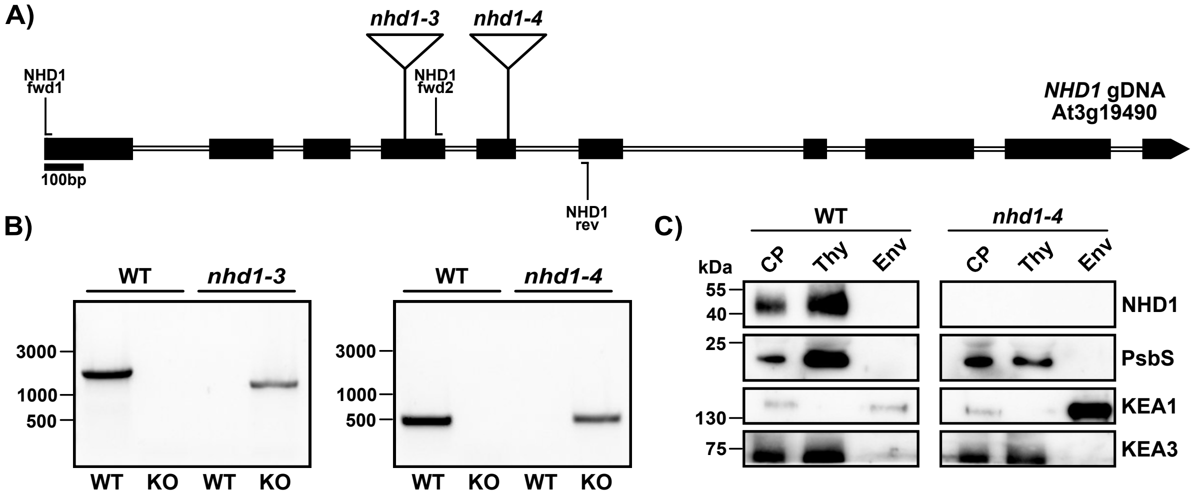


**Fig. S1. NHD1 is a thylakoid membrane protein.** A) Schematic representation of exon and intron structure of the *NHD1* locus, indicating the T-DNA insertions sites of the *nhd1-3* (exon 4) and *nhd1-4* (exon 5) alleles. The T-DNA insertion sites of *nhd1-3* and *nhd1-4* were determined by Sanger sequencing of PCR products amplified using the depicted primers. Primer info is provided in Table S5. B) Representative DNA gel electrophoresis images showing the genotyping PCRs for *nhd1-3* and *nhd1-4*. C) Immunoblot analysis of NHD1 in isolated chloroplast and subplastidial fractions. Chloroplasts (CP), thylakoid (Thy) and envelope (Env) proteins were analyzed. Detection of PsbS, KEA3 (both thylakoid markers) and KEA1 (envelope marker) were used as controls to probe the purity of each fraction.

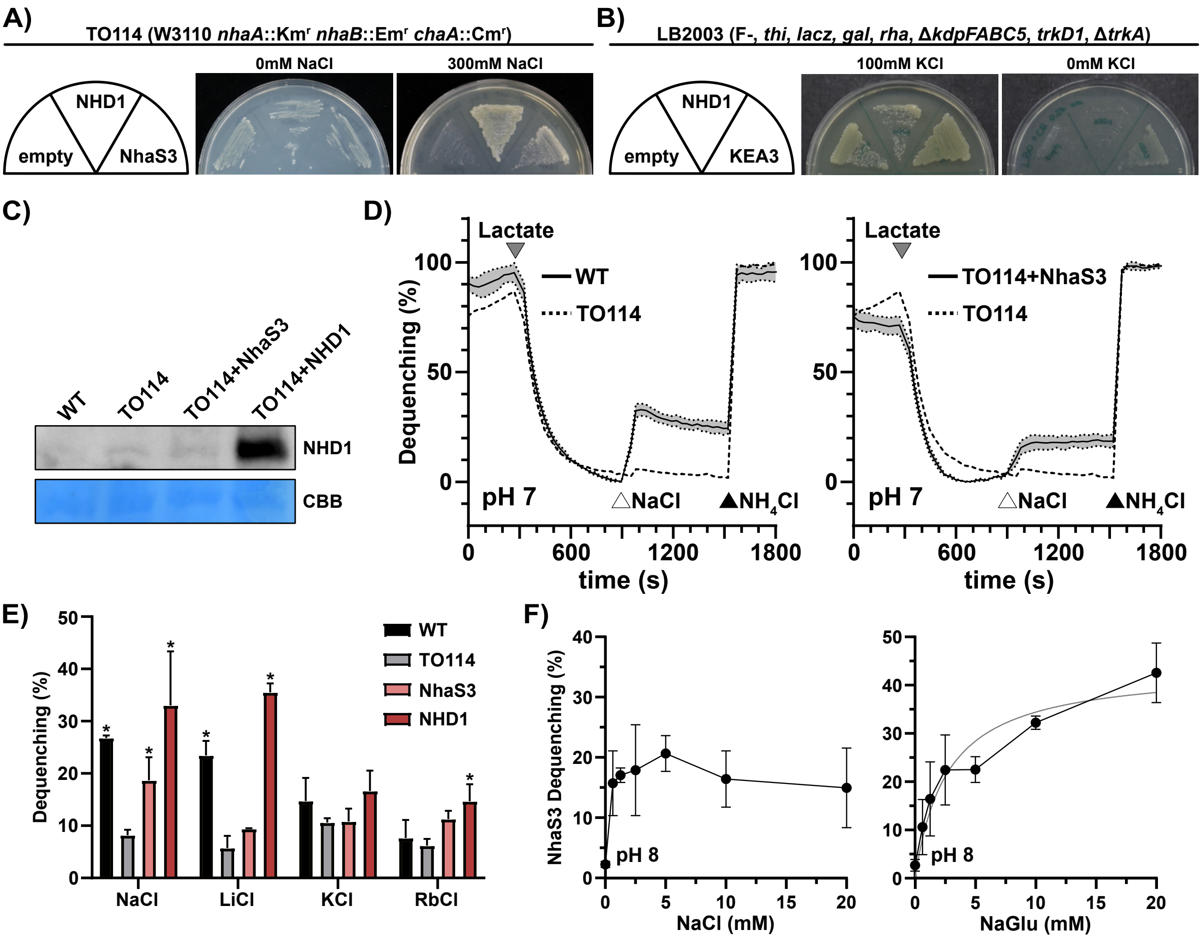


**Fig. S2. NHD1 catalyzes $Na^+/H^+$ exchange.** A) Growth complementation assay of $Na^+$ export-deficient *E. coli* strain (TO114) transformed with *NHD1*, *Synechocystis sp.* PCC 6803 *NhaS3* (positive control), and an empty vector (negative control). Cells were grown in the presence of 300 mM NaCl. B) Growth complementation assay of a $K^+$ import deficient *E. coli* strain (LB2003) expressing NHD1. Transformed cells were grown on LB0 medium and LB0 medium supplemented with 100 mM KCl. Additionally, cells were transformed with *A. thaliana KEA3* (positive control) and the empty vector (negative control). NHD1 expression did not rescue growth of LB2003. C) Immunoblot analysis of NHD1 in inverted vesicles of WT, TO114 and TO114 expressing either NhaS3 or NHD1. D) $Na^+/H^+$ antiporter activity of WT and NhaS3 in inverted membrane vesicles of *E. coli*. Addition of D-Lactate (grey triangle) produces a ΔpH i.e. acridine orange fluorescence is quenched. Antiporter activity is initiated by the addition of NaCl (white triangle). Lastly, ΔpH is dissipated by the addition of $NH_4Cl$ (black triangle). TO114 was used as a negative control (avg. ± SD, n=3). E) Antiporter activity mediated by NHD1 after the addition of NaCl, LiCl, KCl and RbCl in comparison to native $Na^+/H^+$ antiporters of WT, TO114 and *Synechocystis sp.* PCC 6803 NhaS3 (avg. ± SD, n=3). F) Concentration dependent $Na^+/H^+$ antiport of NhaS3 using NaCl (left panel) or Na-D-gluconate (NaGlu) (right panel) at pH 8.0 for NhaS3 containing vesicles (avg. ± SD, n=3). The Michaelis-Menten fit is shown in gray for NaGlu induced antiport.

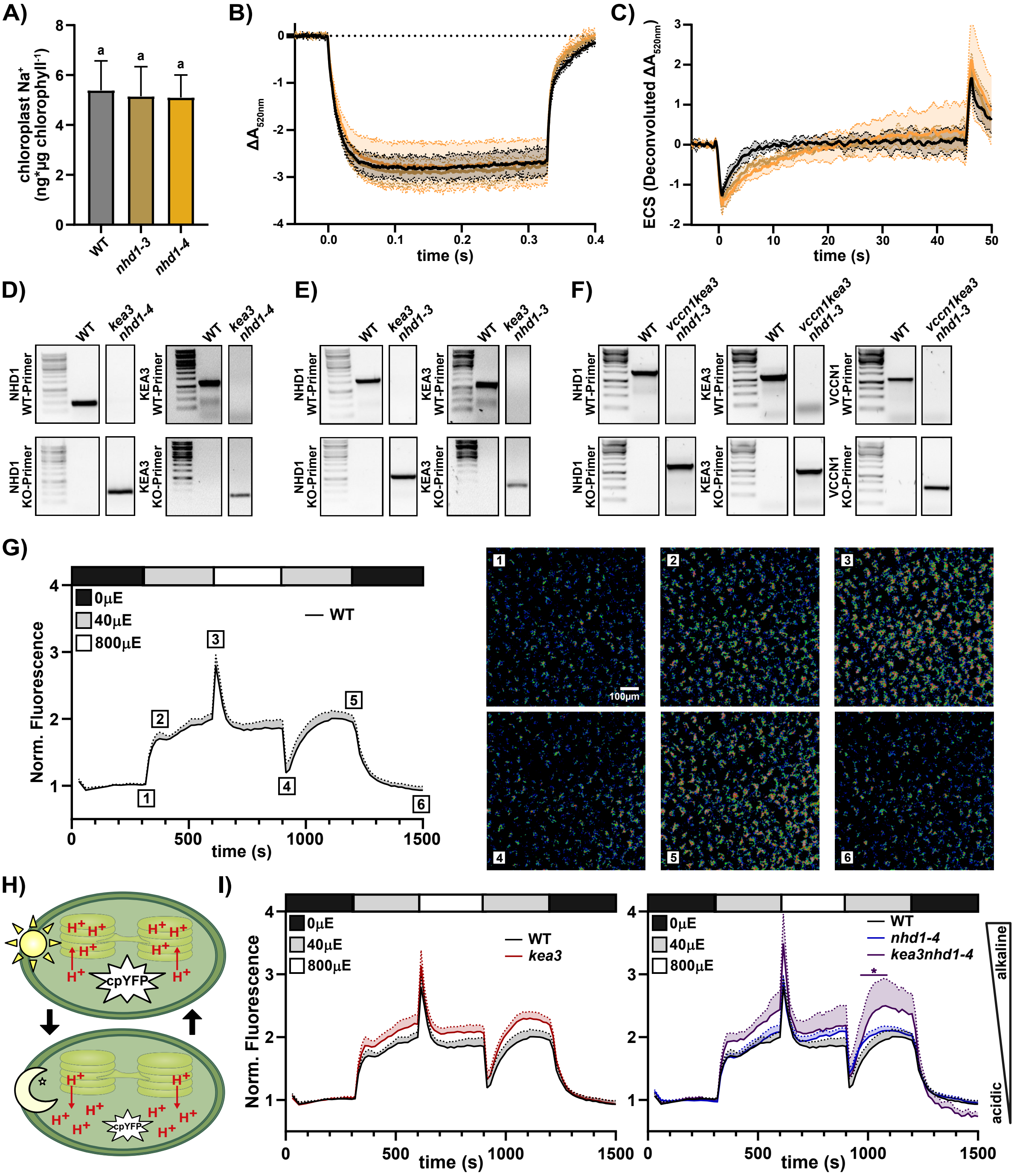


**Fig. S3. NHD1 importance is revealed in the absence of other thylakoid ion transport proteins.** A) $Na^+$ content in isolated chloroplasts, measured by ICP-OES (avg. ± SD, n=6). Statistical significance was determined using a one-way ANOVA (*P*-value < 0.05; letters indicate significant groups). Electrochromic shift (ECS) after light-dark transition in WT and *nhd1* alleles following illumination with 750 µmol photons $m^{-2}$ $s^{-1}$ showing B) 520nm ECS decay during a 330 ms dark interval and C) ECS decay during a long 45 s dark interval. The 520 nm ECS kinetics during the longer dark interval were deconvoluted from the overlapping signals at 505 nm and 535 nm (avg. ± SD, n=3). D), E) and F) Representative DNA gel electrophoresis images showing the genotyping PCRs for *kea3nhd1-3*, *kea3nhd1-4* and *vccn1kea3nhd1-3*. Primer info is provided in Table S5. G) and I) Changes in stromal pH in *A. thaliana* expressing cpYFP in response to changing light conditions. Leaf discs were dark-adapted for 5 min (black box) to establish a baseline, followed by exposure to white light under alternating low light (40 µmol photons $m^{-2}$ $s^{-1}$, gray box), high light (800 µmol photons $m^{-2}$ $s^{-1}$, white box) and low light (40 µmol photons $m^{-2}$ $s^{-1}$, gray box) for 5 min each. Subsequently, measurements were continued for 5 min in the dark. For visual clarity the avg. ± SEM is shown (n≥4). Statistical significance was determined at each time point using a two-way ANOVA (Dunnett's multiple comparisons test; significantly changed comparisons are indicated by an asterisk). G) Intensiometric images of mesophyll expressing cpYFP taken at indicated time points. Scale bar = 100 µm. H) Cartoon of fluorescence dynamics of stroma targeted circular permuted yellow fluorescent protein (cpYFP). Light treatment induces an increase in sensor fluorescence, corresponding to an increase in pH.

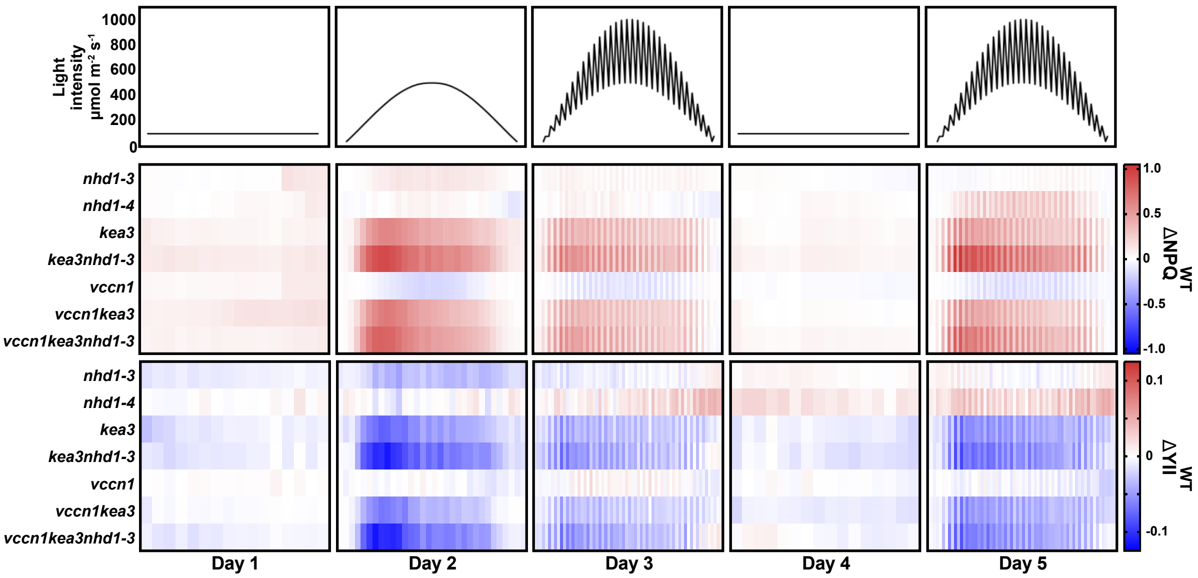


**Fig. S4. Dynamic Environment Photosynthesis Imaging (DEPI) reveals *nhd1* loss-of-function phenotypes under specific light conditions and genetic backgrounds.** Plant photosynthetic performance was monitored over 5 days with different light conditions: flat day (days 1&4), sinusoidal day (day 2), fluctuating day (days 3&5). Shown are NPQ (upper half) and YII (lower half) in the depicted genetic background (avg., n≥10).

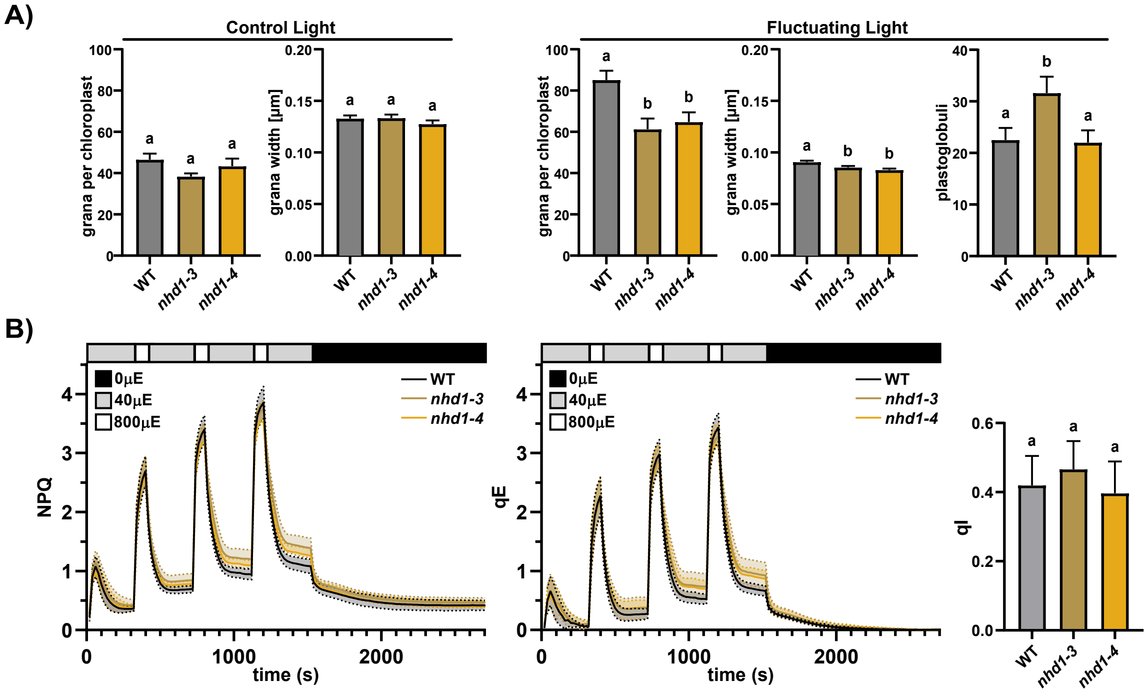


**Fig. S5. NHD1 function is crucial under long-term fluctuating light treatment.** A) Quantification of grana number, grana width and plastoglobuli abundance of WT and *nhd1* loss-of-function mutant chloroplasts following control and fluctuating light treatment, as determined from electron microscopy images (avg. ± SD, n>9). Statistical significance was determined using a one-way ANOVA (*P*-value < 0.05; letters indicate significant groups). B) NPQ and the two NPQ components qE and qI of WT and mutant alleles after growth in fluctuating light. During the measurement, plants were exposed to alternating light conditions consisting of low light (40 µmol photons $m^{-2}$ $s^{-1}$, gray box, 240 s) and high light (800 µmol photons $m^{-2}$ $s^{-1}$, white box, 60 s). Subsequently, measurements were continued in the dark (black box, 15 min) (avg. ± SD, n=9). Statistical significance was determined using a one-way ANOVA (*P*-value < 0.05; letters indicate significant groups).

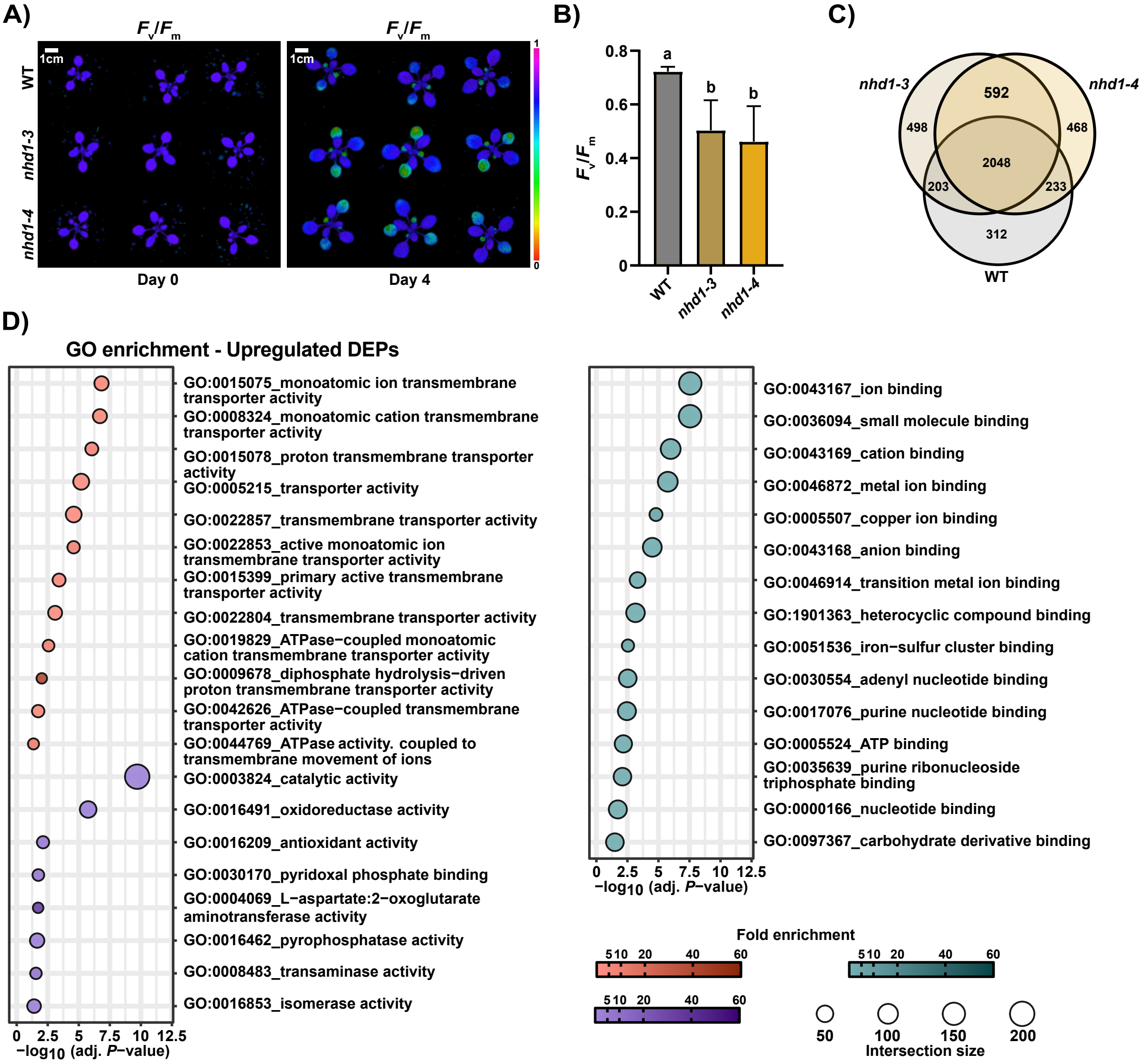


**Fig. S6. Mass spectrometry reveals reorganization of thylakoid proteins in *nhd1* mutants following fluctuating light treatment.** A) Representative $F_v/F_m$ recording after 0 and 4 days of fluctuating light treatment of WT and *nhd1* alleles. B) $F_v/F_m$ of older leaves (leaf 3 and 4) after 4 days of light treatment (avg. ± SD, n=9). Statistical significance was determined using a one-way ANOVA (*P*-value < 0.05; letters indicate significant groups). C) Venn diagram of significantly deregulated proteins in WT and *nhd1* alleles following fluctuating light treatment. 592 proteins are exclusively found to be changed in both *nhd1* mutants. D) Selected significantly enriched GO-terms (Benjamini-Hochberg FDP < 0.05). Circle size corresponds to the number of DEPs within GO-terms, while coloring depicts fold enrichment of manually selected groups. More DEPs included in GO-terms are provided in Table S4.

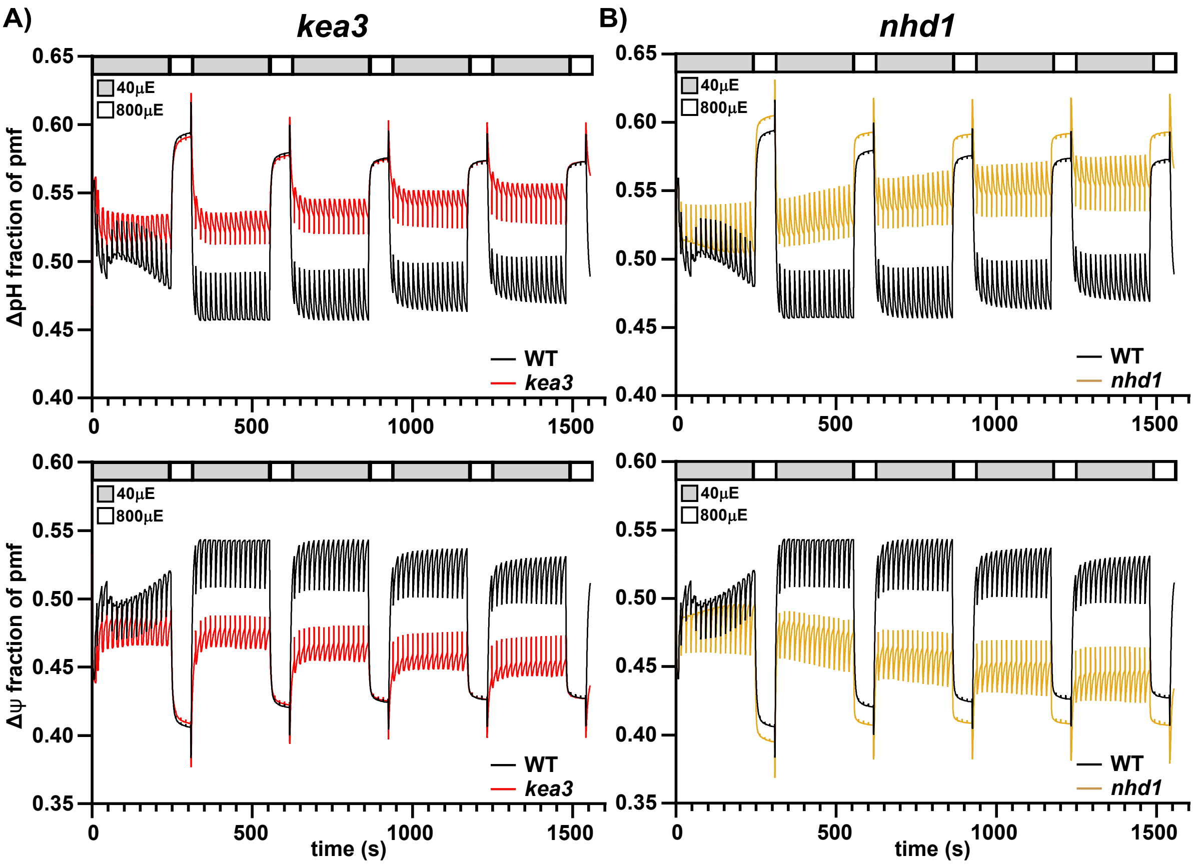


**Fig. S7. Photosynthesis simulation in the presence or absence of thylakoid $K^+/H^+$ and $Na^+/H^+$ antiport, respectively.** A) Extended photosynthesis model based on Ebeling et al., 2026, with $Na^+$ as a third metal ion. Simulation of WT vs. A) *kea3* and B) *nhd1* loss-of-function mutant phenotypes under fluctuating light conditions. Modelled parameters: ΔpH (upper panel) and Δψ (lower panel).
